# CellRegMap: A statistical framework for mapping context-specific regulatory variants using scRNA-seq

**DOI:** 10.1101/2021.09.01.458524

**Authors:** Anna S.E. Cuomo, Tobias Heinen, Danai Vagiaki, Danilo Horta, John C. Marioni, Oliver Stegle

**Affiliations:** European Bioinformatics Institute (EMBL-EBI), Hinxton, Cambridge, UK; Wellcome Sanger Institute, Hinxton, Cambridge, UK; Division of Computational Genomics and Systems Genetics, German Cancer Research Centre (DKFZ), Heidelberg, Germany; European Molecular Biology Laboratory (EMBL), Genome Biology, Heidelberg, Germany; Heidelberg University, Faculty of Mathematics and Computer Science, Heidelberg, Germany; Cancer Research UK, Cambridge Institute, Cambridge, UK

## Abstract

Single cell RNA sequencing (scRNA-seq) enables characterizing the cellular heterogeneity in human tissues. Technological advances have enabled the first population-scale scRNA-seq studies in hundreds of individuals, allowing to assay genetic effects with single-cell resolution. However, existing strategies to perform genetic analyses using scRNA-seq remain based on principles established for bulk RNA-seq. In particular, current methods depend on *a priori* definitions of discrete cell types, and hence cannot assess allelic effects across subtle cell types and cell states. To address this, we propose *Cell Regulatory Map* (CellRegMap), a statistical framework to test for and quantify genetic effects on gene expression in individual cells. CellRegMap provides a principled approach to identify and characterize heterogeneity in allelic effects across cellular contexts of different granularity, including cell subtypes and continuous cell transitions. We validate CellRegMap using simulated data and apply it to two recent studies of differentiating iPSCs, where we uncover a previously underappreciated heterogeneity of genetic effects across cellular contexts. Finally, we identify fine-grained genetic regulation in neuronal subtypes for eQTL that are colocalized with human disease variants.

## Introduction

Seminal population-scale single-cell RNA sequencing (scRNA-seq) studies have demonstrated the feasibility to map expression quantitative trait loci (eQTL) using scRNA-seq as a readout. These studies have successfully replicated eQTL that had previously been discovered using bulk RNA-seq profiles^1,2^, and more importantly, demonstrated increased resolution by mapping eQTL across individual cell types that are captured by scRNA-seq^2–4^.

Despite the scope of these novel opportunities posed by using scRNA-seq for genetic mapping, existing strategies to analyse the resulting data remain largely based on principles that were originally devised for bulk RNA-seq profiling. For example, while established multi-tissue eQTL methods (e.g., refs. ^5–13^) remain in principle applicable to identify eQTL specific to individual cell types, these methods require discretization of the single-cell profiles into distinct cell clusters *a priori* to quantify gene expression. Consequently, these approaches do not fully leverage the resolution provided by single-cell data, potentially failing to detect changes in allelic regulation across subtle cell subtypes. Discretization of single transcriptome profiles into discrete cell clusters can also be limiting in settings where cell states change in a continuous manner, as for example observed across developmental time course or cellular differentiation. Additionally, even seemingly discrete cell types may share common axes of heterogeneity, e.g. due to cell-intrinsic factors such as the cell cycle, thus motivating to jointly analyse genetic effects across multiple cell states in order to capture all of these dimensions.

Here, we propose Cellular Regulatory Map (CellRegMap), a framework for mapping regulatory variants in an unbiased manner across cell types and cell states as obtained from the scRNA-seq profiles. CellRegMap does not require any discretization of cells into cell types, nor is it required to annotate or define specific cell states *a priori*. Instead, the model leverages a multi-dimensional cell state manifold estimated from single-cell transcriptome profiles to define *cellular contexts* in a continuous and unbiased manner. CellRegMap then allows to test for and characterize interaction effects between genetic variants and cellular context on gene expression traits (**Fig. 1**). We validate CellRegMap using simulated data, and apply the model to two recent single-cell genetics studies^1,3^, where we demonstrate increased power to detect interactions, and identify regulatory modules of eQTL that are active in the same cellular contexts. Finally, we demonstrate the relevance of cell-context interactions to fine-map colocalization events with human disease variants.

**Figure 1.**
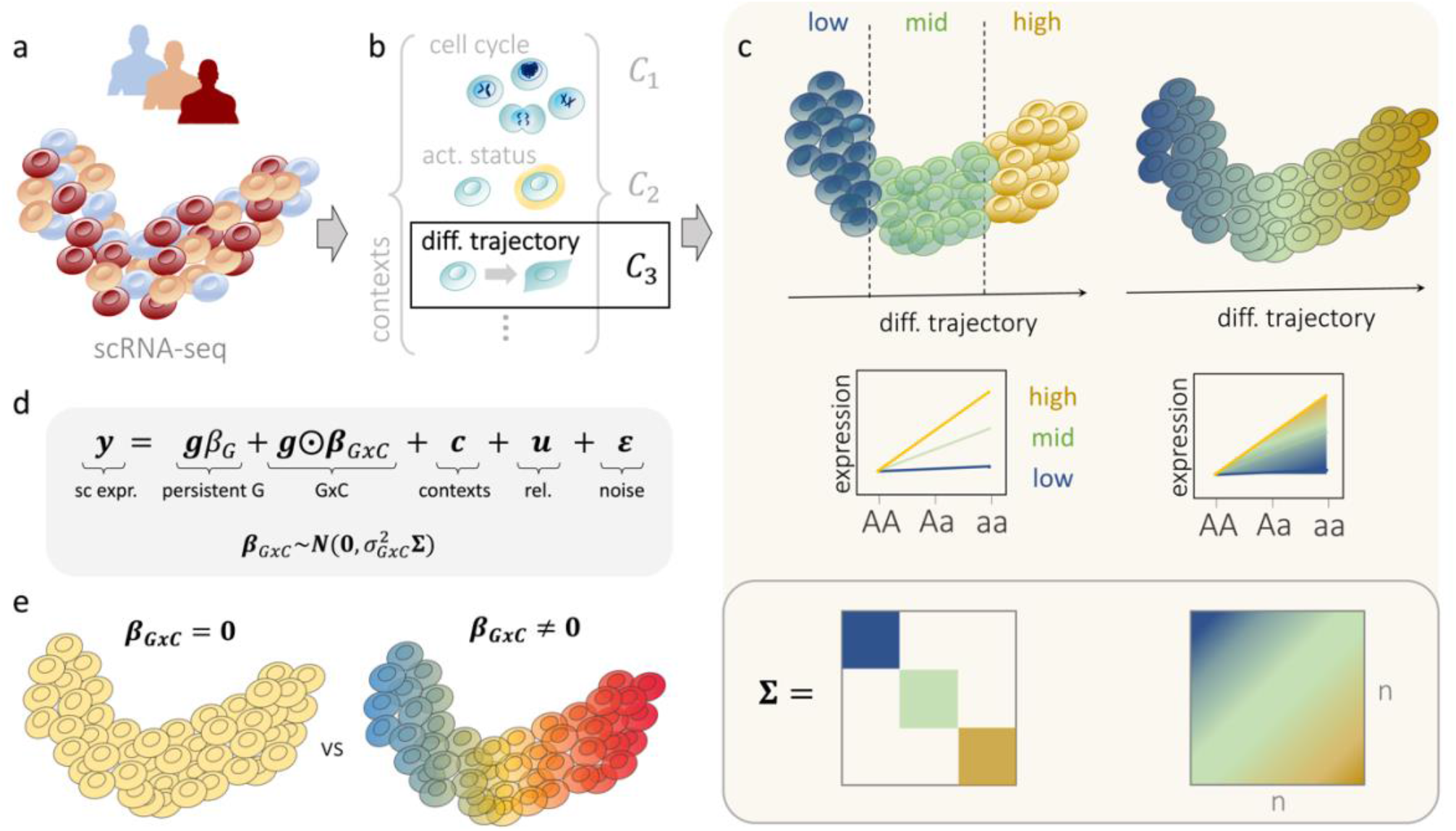
CellRegMap overview and concept. **(a, b)** Established workflows based on principal component analysis or factor analysis applied to scRNA-seq can be used to both estimate cellular manifolds **(a)** and to uncover individual factors that capture different cellular contexts **(b)**. In addition to capturing major cell types, these factors can also explain subtle subtypes, as well as cell-type independent variation, such as the cell cycle and other cell-intrinsic factors. These cellular contexts can represent both discrete or continuous cell state transitions, including cellular differentiation. **(c)** Example of a genotype-context (GxC) interaction across a cellular differentiation context. Left: Established analysis strategies that require discretization into discrete cell clusters (low, mid, high), whereas CellRegMap enables assaying allelic effects as a function of the continuous differentiation context (right). Top panel: cellular manifold with colour denoting allelic effects, either estimated in discrete cell populations (left) or in continuous fashion using CellRegMap (right). Middle panel: Alternative representation of allelic effects for different genotype groups, again considering a discrete (left) or continuous approach (right). Bottom panel: Encoding of discrete cell types (left) and continuous gradients using a cellular context covariance matrix in CellRegMap. (**d**) The CellRegMap model can be cast as linear mixed model, where single-cell gene expression values of a given gene are modelled as a function of a persistent genetic effect, GxC genotype-context interactions, additive effects of cellular context, relatedness and residual noise. GxC interactions are modelled by treating allelic effect size estimates in individual cells (*β*_*GxC*_) as random variable with prior covariance *Σ* (**c**). (**e**) CellRegMap allows to test for heterogeneous genetic effects across cells due to GxC at a given locus for a given gene (testing *β*_*GxC*_ = 0 vs *β*_*GxC*_ ≠ 0).

## Results

CellRegMap generalizes the classical linear interaction model for genotype-environment interactions^2,14^ and allows to test for interactions between genotype and both discrete and continuous cellular context. Briefly, CellRegMap take a cellular context covariance estimated from scRNA-seq as input. This covariance can be derived using existing workflows, including factor analysis (e.g., multi-omics factor analysis, MOFA^15^) or principal component analysis (PCA; **Fig. 1a,b**). CellRegMap incorporates the estimated cellular context covariance to account for interaction effects within the linear mixed model (LMM) framework^16–19^ (**Fig. 1c,d**). Briefly, in addition to a conventional persistent genetic effect, CellRegMap accounts for GxC interactions by modelling heterogeneous genetic effects in individual cells as random effects (**Methods, Fig. 1d**). CellRegMap builds on and extends StructLMM, an LMM-based method to assess genotype-environment interactions in population cohorts^20^. In particular, CellRegMap includes an additional random effect component to account for relatedness between samples, thereby accounting for the repeat structure in single-cell analyses. This is required because typically multiple cells are sampled from the same individuals (**Methods**).

More formally, CellRegMap models the single-cell expression profile of a given gene (across a total of N cells from multiple individuals; *y*) as a sum of a conventional -persistent-genetic effect (G), interactions with cellular context (GxC), additive contributions from cell context (C), a relatedness component (rel.) and residual noise (**Fig. 1d**). GxC interactions are modelled as an element-wise product between the expanded genotype vector **g** at a given locus and a GxC effect size vector 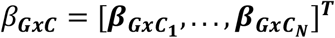, which correspond to allelic effect sizes in individual cells. This vector follows a multivariate normal distribution, 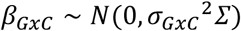. Depending on the parametrization of the cell-context covariance *Σ*, CellRegMap can be setup to account for interactions with different cellular contexts, including discrete and related cell types, as well as continuous cell state transitions (**Fig. 1c,d, Methods**). The same covariance is also used to account for additive effects of cellular context on expression, i.e., 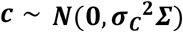. To account for the repeat structure caused by sampling multiple cells from the same individual, CellRegMap includes an additional relatedness component, which is parametrized as a product covariance between a conventional kinship covariance ***K*** and the cell context covariance, i.e., 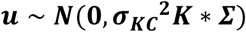 (see **Methods**). This component ensures that the model retains calibration when sampling multiple cells from the same individual. Finally, the model assumed Gaussian distributed and independently and identically distributed residual noise, i.e. 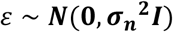 **(Fig. 1e)**.

We propose a score test to identify gene-loci pairs with significant G×C effects (testing *β*_*GxC*_≠ **0, Fig. 1f**), which generalizes the approach in ref. ^20^. Additionally, CellRegMap can be used to characterize GxC effects of eQTL by estimating the allelic effect for individual cells *β*_*GxC*_, which can be used to identify specific cell populations with increased or decreased genetic effects (**Fig. 1c, Methods**). The model is implemented in efficient open-source software, which leverages low-rank representations and factorizations of the resulting total covariance, after marginalizing the random effect components (**Methods**). As a result, the computational complexity of CellRegMap scales linearly in the number of cells (**Supp. Fig. 1**.**1**).

## Model validation using simulated data

Initially, we considered simulated data to validate the calibration of CellRegMap and to assess statistical power of the model. We simulated single-cell expression profiles by sampling from the generative model of CellRegMap (**Methods**), using a cell context covariance *Σ* derived from the leading 20 principal components of a real scRNA-seq dataset (from ref. ^1^; see below).

We confirmed statistical calibration of CellRegMap, both when considering expression phenotypes generated from a null without simulated genetic effect (**Supplementary Fig. 2**.**1**), or when simulating from a persistent effect model without GxC interactions (**Fig. 2a,b**, **Supplementary Fig. 2**.**1; Methods**). We also compared CellRegMap to a reduced model that does not include the relatedness component (which is similar to the existing StructLMM^20^ model), thereby confirming that the relatedness component is required to retain calibrated p-values when multiple cells are assayed from each individual (**Fig. 2a,b**, **Supplementary Fig. 2**.**1**).

**Figure 2.**
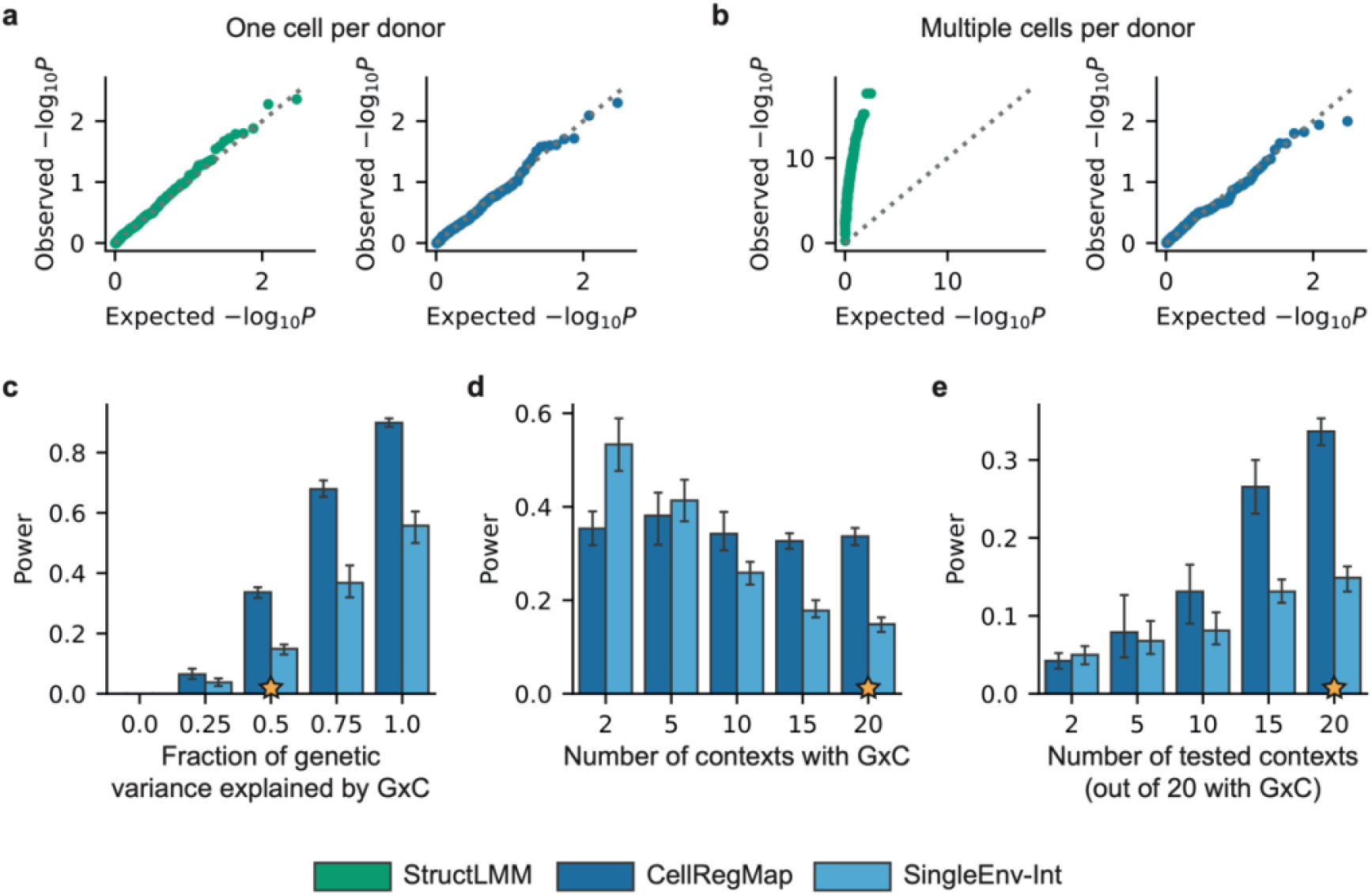
CellRegMap validation using simulated data. **(a, b):** Assessment of statistical calibration of the GxC interaction test based on 200 variant-gene pairs simulated from the null (persistent genetic effect, no GxC interaction; Methods), either simulating a single cell per individual (**a**) or simulating multiple cells for each individual (50 cells per individual; **b**). Shown are QQ plots of expected versus observed negative log p-values for StructLMM and CellRegMap. Whereas both StructLMM and CellRegMap are calibrated if a single cell is sampled per individual, the relevant setting of multiple cells per individual requires the additional relatedness component in CellRegMap to retain calibrated results. **(c-e):** Assessment of power to detect simulated GxC effects. Shown are results from CellRegMap and a single-environment interaction test (SingleEnv-Int) with the same relatedness component as used in CellRegMap (Bonferroni-adjusted for the number of cell contexts; Methods). **(c)** Power as a function of the fraction of genetic variance explained by GxC. **(d)** Power as a function of the number of active contexts (contexts with nonzero GxC contribution), when testing 20 cellular contexts. **(e)** Power as a function of the number of tested contexts (out of 20 all contributing to GxC). Results presented are based on 250 stimulated genes, with bars hight corresponding to average power and error bars to standard deviations estimated across 10 repeat experiments. Stars denote the default value of that parameter when other parameters are being varied.

Next, we conducted experiments to assess the statistical power of CellRegMap for identifying loci with simulated GxC effects. We simulated single-cell expression profiles with variable fractions of genetic variance explained by GxC (**Fig. 2c; Methods**). For comparison, we also considered a conventional linear interaction test (similar to the approach in ref. ^14^) that assesses a linear interaction with individual cellular contexts (SingleEnv-Int; adjusted for multiple testing across factors using Bonferroni; **Methods**), but using otherwise the same random effect components to account for additive effects of context and relatedness employed in CellRegMap (**Methods**). The power of both tests increased as the fraction of the genetic effect explained by GxC increases, noting that CellRegMap was substantially better powered than the SingleEnv-Int test (**Fig. 2c, Supplementary Fig. 2**.**2**). As a second parameter, we varied the number of cellular contexts that are simulated to contribute to G×C (out of 20 included in both tests). The results of this analysis show that CellRegMap outperformed the corresponding SingleEnv-Int GxC test when larger numbers of cellular contexts contribute to GxC (**Fig. 2d, Supplementary Fig. 2**.**2**). We also varied the number of cellular contexts tested in the model, when all 20 contribute to GxC (**Fig. 2e, Supplementary Fig. 2**.**2**). Finally, we repeated the simulations using a negative binomial model to simulate observation noise, as expected in real scRNA-seq data, and we assessed the effect of sampling variable numbers of cells per individual, consistently observing calibrated test statistics and power benefits of CellRegMap (**Supplementary Fig. 2**.**1, 2**.**2; Methods)**. Taken together, these results demonstrate power advantages and robustness of CellRegMap, compared to existing methods, particularly when multiple cellular contexts contribute to GxC.

## Application to a continuous trajectory of iPS cells differentiating towards definitive endoderm

Next, we applied our model to a recent single-cell RNA-seq dataset of differentiating induced pluripotent stem cells (iPSCs) that spans 125 genetically diverse individuals^1^. Briefly, a total of ∼30,000 cells were captured at four time points of iPSC differentiation (day0: iPSCs, day1, day2 and day3 of differentiation towards definitive endoderm; **Fig. 3b**), using the SMART-Seq2^21^ protocol. As expected, cell differentiation is the dominant cellular context in this study, and hence this dataset is an ideal test case to assess the ability of CellRegMap to identify continuous changes of allelic effects across a cellular trajectory.

**Figure 3.**
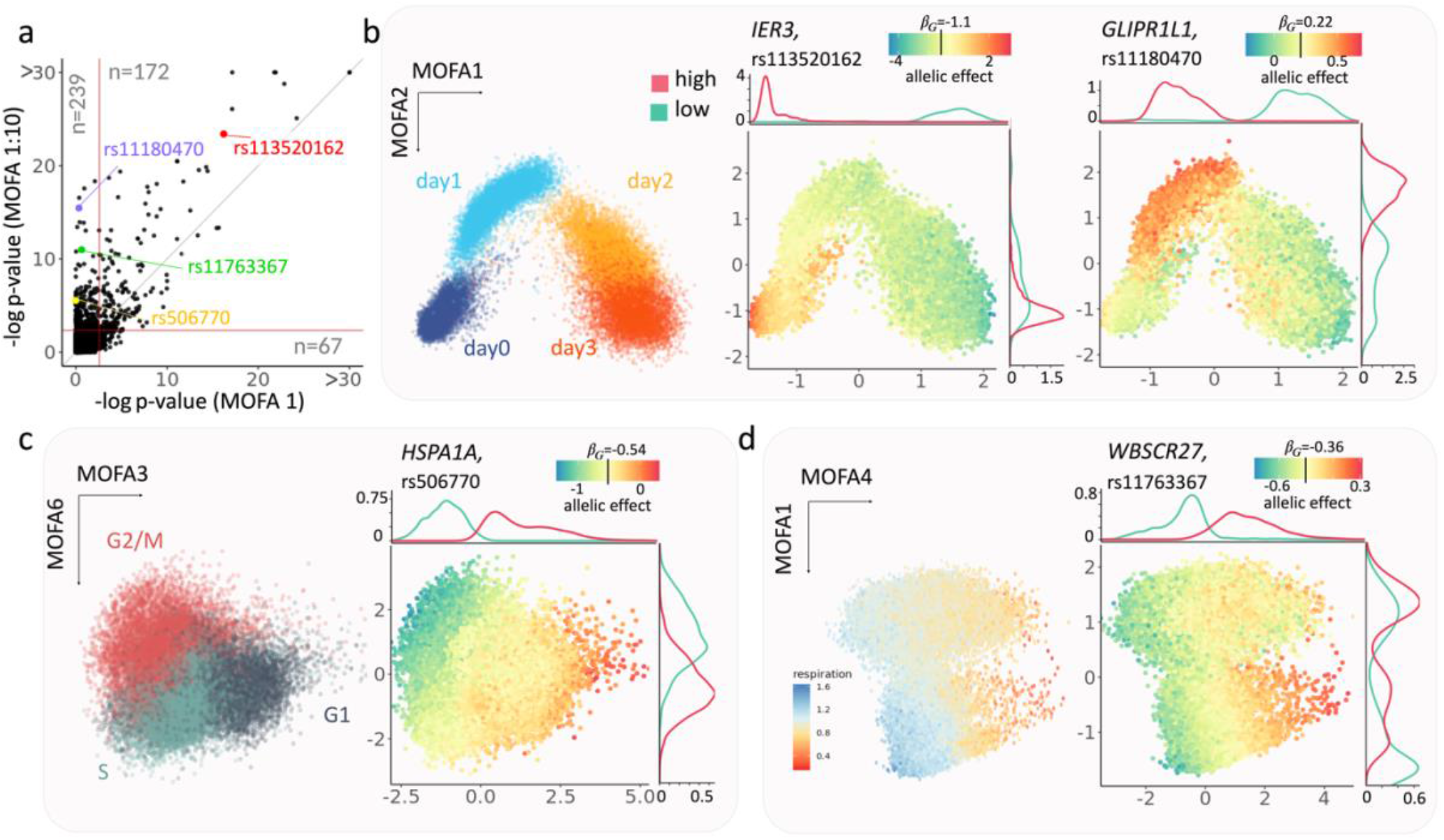
Application of CellRegMap to iPSCs differentiating towards definitive endoderm. **(a)** Comparison of the CellRegMap GxC interaction test, either considering the first MOFA factor to define the cell context covariance (MOFA 1, x axis) versus using the leading 10 cellular factors (MOFA 1:10, y axis). Shown is a scatterplot of negative log p-values obtained from the respective tests applied to 4,470 eQTL variants and genes. Horizontal and vertical lines denote the FDR<5% significance threshold (Benjamin-Hochberg adjusted). Shown in each quadrant is the number of eQTL with evidence for a GxC effect (e.g., 239 GxC effects are exclusively detected by the model based on 10 factors; FDR<5%). (**b-d)** Representative examples of eQTL with GxC interaction. **(b)** Left: scatter plot of the first two MOFA factors (capturing cell differentiation as context) with colour denoting the time point of collection (day 0,1,2 & 3 of endoderm differentiation); middle: Identical scatter plot with colour encoding the estimated allelic effect for the eQTL variant rs113520162 for the gene *IER3*; right: allelic effect for the eQTL at rs11180470 for the gene *GLIPR1L1*. Shown are total allelic effects (*β*_*G*_ + *β*_*GxC*_) for individual cells. The allelic effect size colour bar is cantered on the persistent genetic effect (*β*_*G*_). Panels on the top and right display marginal densities of cells that either have increased (high, red) or decreased (low, cyan) allelic effects (corresponding to the bottom and top 10% quantiles respectively). Whereas the GxC effect for the eQTL for *IER3* is primarily explained by the first MOFA component, the GxC effect for *GLIPR1L1* is captured by the combination of the first two MOFA factors. **(c)** Analogous presentation as in **b**, displaying a scatter plot between MOFA factors 3 and 6 with cells coloured by alternative annotations. Left: inferred cell cycle phase (Methods); Right: allelic effects for an eQTL at rs506770 for *HSPA1A* (yellow). **(d)** As in **b,c** scatter plot of MOFA factors 4 and 1. Left: cells coloured by cellular respiration (Methods); Right: allelic effects for the eQTL at rs11763367 for *WBSCR27* (green).

We used MOFA^15^ to infer latent factors that explain variation in cellular contexts in the data, which captured both differences in major cell types across the differentiation trajectory, but also more subtle cell states. For example, the first factor (MOFA 1) primarily explained the differentiation axis, with cells transitioning between a pluripotent state and the definitive endoderm fate. Higher order factors captured other cellular contexts, including cell cycle phase (MOFA 3 and 6), respiration (MOFA 4) and others (**Supplementary Fig. 3**.**1; Methods**).

We applied CellRegMap to test for GxC effects at 4,470 eQTL variant / gene pairs that were previously identified in the primary analysis of the dataset using a conventional eQTL mapping workflow that does not account for GxC interactions^1^. We compared CellRegMap when only using the first MOFA factor to define the cell context covariance, which is similar to the approach taken in the primary analysis^1^, to a model that leverages the information contained in the leading 10 MOFA factors. The model with 10 components yielded a substantially larger number GxC effects (322 versus 183, FDR<5%; **Fig. 3a, Supplementary Table 1**), indicating that despite cell differentiation being the major driver of expression variation, other more subtle cellular states also manifest in GxC interactions on gene expression.

Next, we set out to characterize specific cellular contexts that are associated with the identified GxC interactions. We used CellRegMap to estimate the GxC allelic effect component in each cell, thereby recovering the continuous landscape of the GxC component of genetic effects across the cell-context manifold (**Methods**). This analysis identified a range of allelic patterns, including GxC effects that are primarily governed by cellular differentiation but also more complex patterns that involve multiple cellular contexts and higher-order cellular factors. For example, the eQTL variant rs113520162 for *IER3* had a GxC effect that reflects variation across cell differentiation explained by the first MOFA component (**Fig. 3b, middle**). Other eQTL, such as rs11180470 for *GLIPR1L1*, had GxC effects that were associated with two MOFA factors (**Fig. 3b, right**). More generally, we observed that higher order MOFA components capture changes in cellular contexts beyond cellular differentiation, including the cell cycle (**Fig. 3c**), cellular respiration (**Fig. 3d**), and others (**Supplementary Fig. 3**.**2**). Collectively these results illustrate how CellRegMap can be used to uncover different cellular contexts that manifest in GxC interactions.

## Application to iPSC-derived dopaminergic neurons

Next, we applied CellRegMap to a single-cell dataset of 215 iPS cell lines that were assayed at three stages of differentiation towards dopaminergic neurons^3^ (11, 30 and 52 days of differentiation) using the 10X Genomics technology (3’ kit^22^), as well as a stress condition at the most differentiated time point. These data feature prominent discrete cell states rather than continuous changes, thus providing a complementary use case.

To assess whether CellRegMap can identify GxC effects associated with finer grained neuronal subtypes, we considered 147,801 cells that were annotated as dopaminergic neurons in the primary analysis of this dataset (based on marker genes^3^). This selection included cells collected at three of the four time points and conditions: young neurons (at day 30 of iPSC differentiation), mature neurons (day 52) and mature neurons followed by rotenone treatment (day 52 ROT). A t-SNE embedding of these cells identified discrete cell populations that reflect the combination of differentiation stage and stimulus (**Fig. 4a**). Our hypothesis is that while it is expected that regulatory variants can be specific to these major sub populations, there could also be GxC effects that are more granular due to cellular contexts that capture subpopulations within these clusters, or that capture shared cellular contexts that are present across these clusters. To mitigate the sparsity of 10X sequencing data compared to SMART-Seq2, we aggregated the read count information into pseudocells (similar to approaches described in refs. ^23,24^; resulting in 17 cells on average, 8,648 pseudocells in total, **Supplementary Fig. 4**.**1; Methods**). We again considered the leading 10 MOFA components to define the cell context covariance for analysis using CellRegMap.

**Figure 4.**
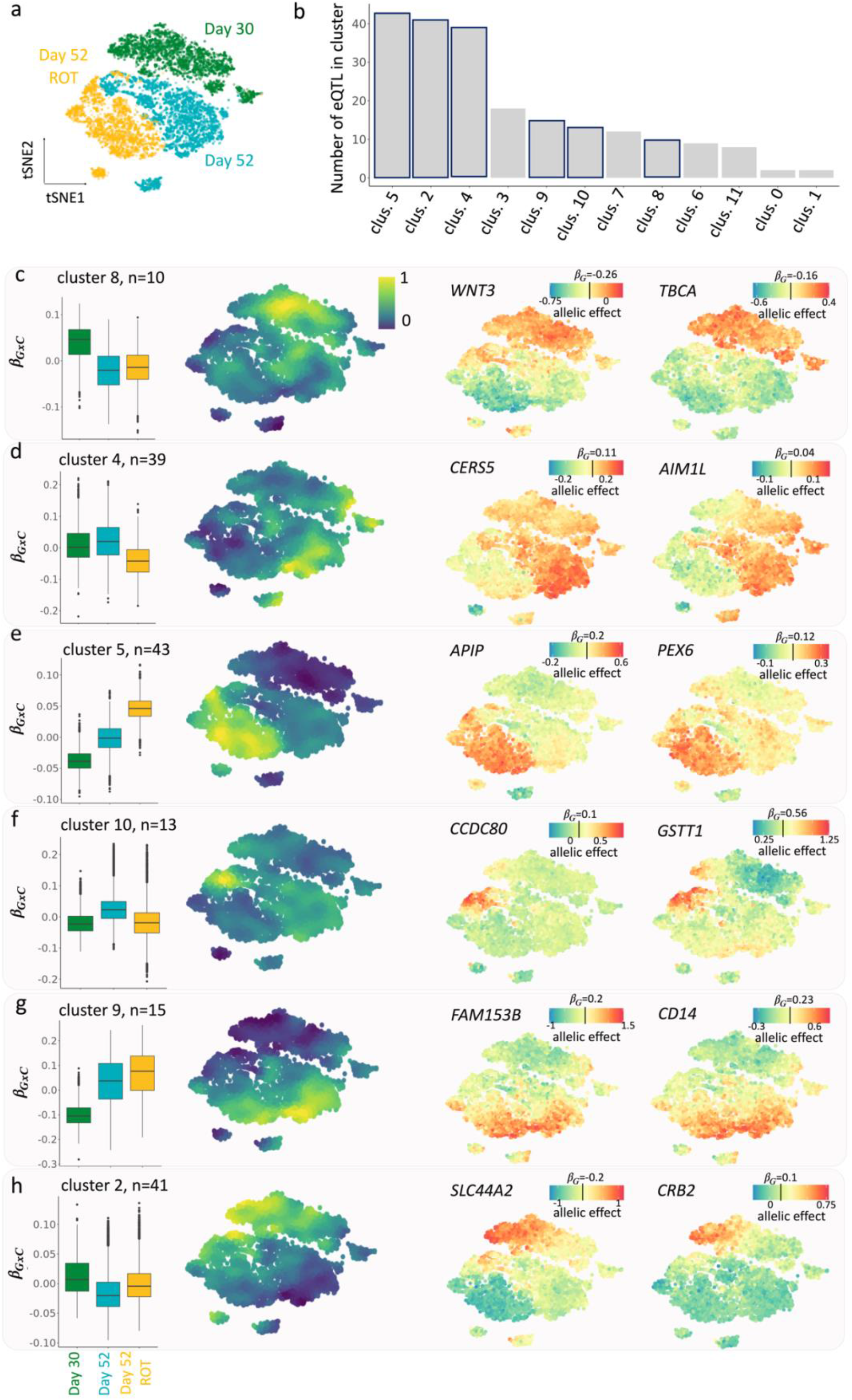
CellRegMap identifies fine-grained regulatory modules in dopaminergic neurons. **(a)** Overview of the cell subpopulations: tSNE plot of 8,648 pseudocells (Methods), highlighting three major populations of dopaminergic neurons: young neurons (day 30 of iPSC differentiation), more mature neurons (day 52) and rotenone-treated day 52 dopaminergic neurons (day 52 ROT). **(b-h)** Results from clustering of allelic effect profiles of GxC interactions based on relative allelic effect size estimates. **(b)** Barplots indicating the number of eQTL with GxC effects assigned to each cluster. Highlighted are the six representative clusters that are displayed in subsequent panels. **(c-h)** For each of 6 representative clusters, from left to right: box plot of the distribution of the relative GxC effect sizes estimates for cells in each of the three major population, manifold of consensus relative GxC effect sizes estimates for each cluster **a**; example eQTLs with allelic effect size estimates across the cell manifold as in **a** with colour denoting total allelic effects (*β*_*G*_ + *β*_*GxC*_); the colour bar is centred on the persistent genetic effect size estimate for each eQTL (*β*_*G*_).

We tested for GxC effects at 1,374 SNP-gene pairs identified as eQTL in at least one of the three discrete cell populations in the primary analysis of the data^3^ (FDR<5%; **Methods**). This identified 213 eQTL with evidence for GxC interactions (FDR<5%, **Methods, Supplementary Table 2**). We again used estimated allelic effects in single cells to interpolate the landscape of GxC effects for each of the eQTL with a significant GxC effect. To identify principal patterns of genetic regulation across cell contexts, we adapted a clustering approach originally designed for spatial transcriptomics data to group the identified allelic effect patterns across the cell context manifold (implemented in SpatialDE^25^, **Supplementary Fig. 4**.**2a**). This identified 17 clusters with distinct GxC effect profiles (**Fig. 4b**). In each of these clusters, we annotated the subpopulation of cells with the largest absolute GxC effects by identifying genes with covarying expression patterns. Briefly, for each cluster we ranked genes by the correlation between their single-cell expression profiles and the pattern of absolute GxC allelic effects. Based on this gene ranking we then assessed enrichments of known pathways (over-representation analysis using a hypergeometric test and annotations from GO, Reactome, KEGG, HPO and others), as well as using literature curated marker genes of dopaminergic neurons (see **Methods** for details, **Supplementary Fig. 4**.**2b**).

Some of the clusters primarily captured genetic effects that were specific to the three major cell populations. For example, cluster 8 captured eQTL that are primarily active in the day 30 population, cluster 4 eQTL are primarily active in day 52 cells, and cluster 5 captures effects specific to the rotenone-treated day 52 cell population (**Fig. 4c-e**). Gene enrichment analysis of these clusters yielded processes that are consistent with the expected function of the corresponding cell populations, such as response to oxidative stress (GO: 006979) for cluster 5 (**Methods, Supplementary Fig. 4**.**2**).

Beyond these expected patterns of GxC effects, other eQTL had interaction effects that were explained by clusters that show continuous changes of allelic effects across developmental time, or that are specific to more fine-grained sub populations (**Fig. 4f-h**). For example, cluster 10 captured eQTL that are active in common subpopulation of day 52 treated and untreated cells (**Fig. 4f**). Functional enrichment analysis linked this cluster to processes related to exocytosis and neurotransmitter transport through synapsis, suggesting an association with neurons that are actively transmitting cell-cell information (**Supplementary Fig. 4**.**2**). Clusters 2 and 9 exhibit GxC effects with opposing directions, with cluster 2 being associated with increased genetic effects and cluster 9 with decreased effects. Cluster 9 eQTL have increasing absolute effect sizes in more mature neurons, regardless of the stimulation status. Enrichment of this cluster highlights neuronal-specific features such as synaptic signalling (GO:0099536, **Fig. 4g, Supplementary Fig. 4**.**2**). Cluster 2, on the other hand, is specific to a subpopulation of day 30 cells (**Fig. 4h**) that corresponds to less mature dopaminergic neurons, as evident by continuous gradients of canonical dopaminergic neuronal markers (**Supplementary Fig. 4**.**2, Methods**).

Finally, we considered a subset of 94 eQTL with evidence for statistical co-localization with neuronal phenotypes and human disease traits^3^ (**Methods**). Out of these, 14 eQTL had significant GxC interactions. For example, the eQTL variant rs1972183 for SLC35E2 has a GxC effect explained by cluster 4 and is colocalized with a GWAS variant for sleeplessness and insomnia in the subpopulation of day 52 untreated cells. CellRegMap allowed for fine-mapping a specific sub-populations within this cluster with elevated allelic effect sizes (**Supplementary Fig. 5**.**1a)**. We used allelic effect size estimates to mark cells in the top and bottom and bottom 30% quantiles ranked by the absolute GxC effects, and applied a classical *cis* eQTL mapping workflow based on expression estiamtes derived from scRNA-seq counts across cells in the respective quantiles^26^. This analysis confirmed the expected difference in effect sizes (**Fig. 5e,f**), but also highlighted subtle differences in the *cis* eQTL mapping profile for each of these traits (**Fig. 5d**). Notably, the trait associated with the top quantile of increased allelic effects also yielded higher evidence for co-localization with the disease GWAS signal (**Supplementary Fig. 5**.**2**).

**Figure 5.**
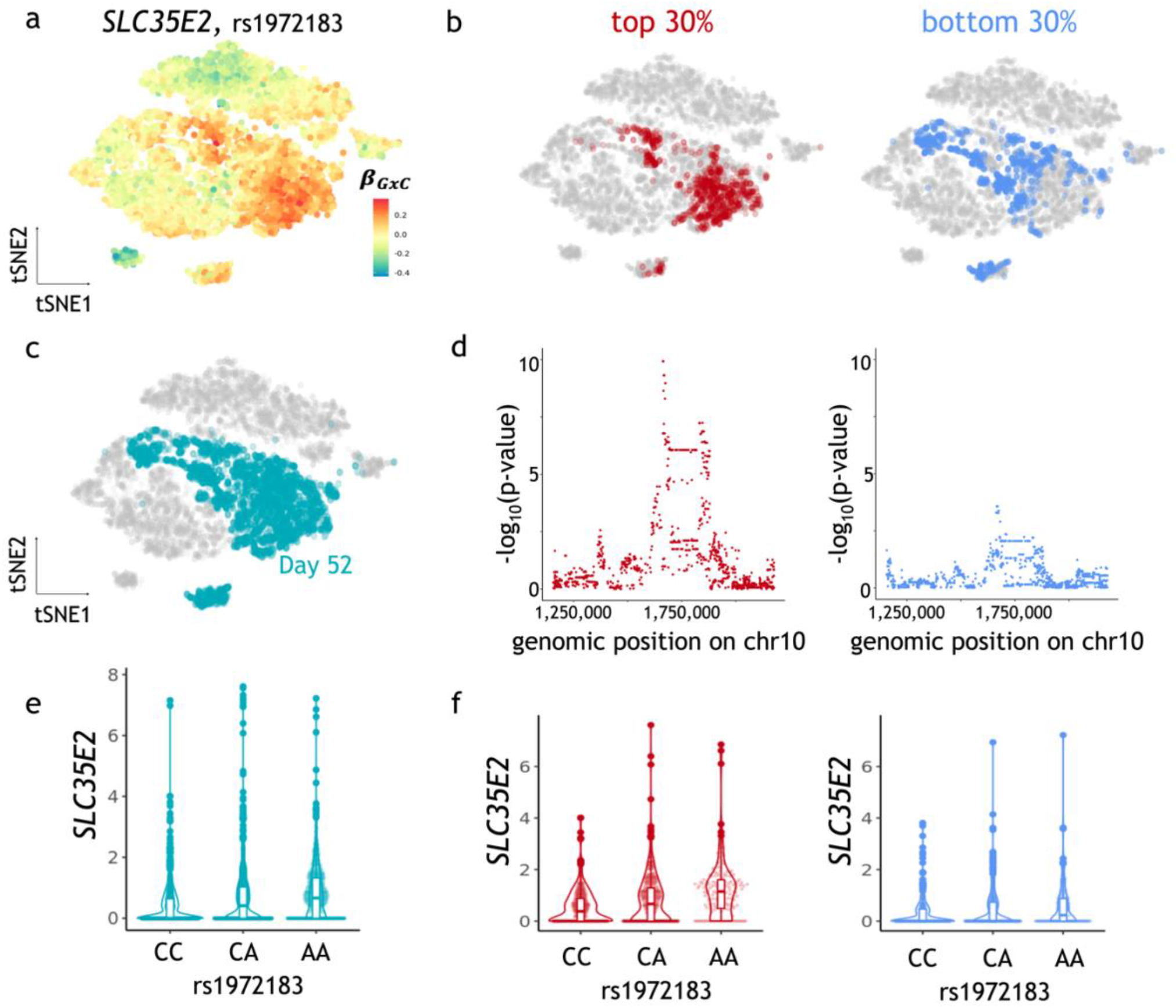
CellRegMap allows to fine-map sub populations of cells linked to human disease variants. **(a)** Allelic effect size estimates for the rs1972183 on *SLC35E2*. Shown is a scatter plot of tSNE coordinates with color denoting estimated GxC allelic effects (*β*_*GxC*_). **(b-f)** *SLC35E2-*eQTL results obtained from a conventional eQTL mapping workflow (Methods), using the CellRegMap output to select alternative cell populations to estimate expression phenotypes. **(b-c)** tSNE plots as in **a**, with colour indicating alternative selected subpopulations of day 52 untreated cells. **(b)** Top and bottom 30% quantiles of cells ranked by the absolute GxC allelic effect. **(c)** Day 52 untreated cells. **(d)** Manhattan plots displaying negative log p-value from a conventional eQTL workflow when using the subpopulations as in **b** to estimate expression phenotypes. Shown are negative log p-vlaues (y-axis) as a function of the genomic position of common variants (x-axis). **(e-f)** Violone plots displaying effect size estimates on *SLC35E2* (y-axis) stratified by genotype at the lead variant rs1972183 (x-axis), either considering all cells for pseudo bulk expression estimation (**e**) or (**f)** considering the subpopulations as in **b, d**.

## Discussion

Here, we presented cellular regulatory map (CellRegMap), a linear mixed model for the identification of context-specific eQTL that is applicable to cellular states derived from scRNA-seq. Critically, CellRegMap overcomes the need to define cellular contexts *a priori* (**Fig. 1**) and instead uses cell manifolds derived from single-cell transcriptome profiles to estimate cellular contexts in an unbiased manner to then test for interaction effects.

Conceptually, CellRegMap is related to and builds on StructLMM, a model that was originally designed to identify genotype-environment interactions in population cohorts^20^. CellRegMap adapts these principles to single-cell genomics, by including an additional relatedness component in the model that accounts for dependencies across cells that are assayed from the same individual. CellRegMap retains calibrated test statistics (**Fig. 2a,b** and **Supplementary Fig. 2**.**1**) and enjoys power benefits compared to conventional fixed-effect interaction tests (**Fig. 2c-e** and **Supplementary Fig. 2**.**2**).

To illustrate the model, we applied CellRegMap to a single-cell dataset of iPS cells from 125 individuals across differentiation towards a definitive endoderm fate^1^. The main source of variation in this dataset is a continuous differentiation signal, which manifests in dynamic eQTL across differentiation. Notably, we also identify eQTL associated with other dimensions of transcriptome variation, including factors associated with cell cycle phase or respiration (**Fig. 3**). As a second-use case, we applied CellRegMap to scRNA-seq data from iPSCs differentiating toward dopaminergic neurons^3^. Our analysis demonstrated that cell-type specific eQTL are not only observed for major subpopulations linked to known cell types, but instead a substantial number is driven by other more subtle variations in cellular context (**Fig. 4**).

An important insight from both use cases is that continuous and subtle allelic regulation, which manifests in GxC in specific subpopulations, is common. Even in seemingly well-defined cell populations, CellRegMap identified heterogeneity in genetic effects that manifests in GxC interactions. These interactions are particularly relevant if they are linked to eQTL with evidence for colocalization with human disease variants. We illustrated this for one disease-linked GxC effect, where CellRegMap allowed to fine-map the specific subpopulation of cells that is primarily responsible for this eQTL signal. Notably, this step does not only enhance the interpretation of most relevant cell populations, but can also yielded more fine-grained *cis* eQTL signals for mapping variants.

Although we demonstrated that CellRegMap is broadly applicable to different datataset and scRNA-seq technologies, the model is not free of limitations. At present, CellRegMap is primarily designed as a tool to annotate known eQTL variants rather than facilitating variant dsicovery. This is analogous to the two-stage strategy for mapping of genotype-environment interactions at known GWAS loci in population cohorts. Such procedures build on the assumption that the persistent genetic effect signal is sufficiently strong to enable discovery. Future extensions of CellRegMap could consider the benefits of accounting for GxC for the purpose of variant discovery itself. A second limitation of the model is that it currently requires appropriate processing steps to provide cell-level or pseudo-cell expression estimates that can be treated as Gaussian distributed traits. Although our results indicate that this approximation is acceptable in practice and retains statistical calibration (**Supplementary Fig. 2**.**2)**, explicit modelling of count data could provide additional power benefits, in particular in the regime of lowly expressed genes. Finally, as datasets grow in size, future developments on the scalability may be warranted. While CellRegMap scales linearly in the number of cells already, the computations required to account for the relatedness component could be prohibitive when analysing very large datasets from thousands of individuals. Datasets of this magnitude could become available through large data-integration efforts, for example via federated analysis envisioned in the single-cell eQTLGen consortium^27^.

## Methods

A complete methods section, including definition of the CellRegModel, simulation strategies and data processing and analyses is available as **Supplementary Methods**.

## Supporting information

Supplementary Figures and Tabels

## Code availability

CellRegMap is available under an open-source license at: https://github.com/limix/CellRegMap/.

Code to reproduce the specific analyses presented here can be accessed under: https://github.com/annacuomo/CellRegMap_analyses/.

## Data availability

The datasets used are accessible at https://zenodo.org/record/3625024 (for the data from ^1^) and https://zenodo.org/record/4651413 (for the data from ^3^).

## Acknowledgements

The authors would like to thank Marc Jan Bonder for helpful discussions. A.C. is supported by an EMBL International PhD Programme (EIPP).

## Author contributions

AC and OS conceived the method. AC and DH implemented the methods. AC performed computational analyses. AC and TH devised the simulation strategy and TH ran the simulation analyses. DV performed some data processing tasks. AC, JM and OS interpreted the results and wrote the manuscript.

