## Supplementary Figures and Tabels for "CellRegMap: A statistical framework for mapping context-specific regulatory variants using scRNA-seq"

Supplementary Information for:

### CellRegMap: A statistical framework for mapping context-specific regulatory variants using single cell RNA-sequencing

Cuomo *et al.*

#### Table of Contents

***Supplementary Tables*** ........................................................................................................ ***3***

***Supplementary Figures***....................................................................................................... ***4***

### Supplementary Tables

**Supplementary Table 1:** List of CellRegMap results for iPSC differentiation toward endoderm (from Cuomo et al, 2020), FDR < 5%

**Supplementary Table 2:** List of CellRegMap results for iPSC differentiation toward dopaminergic neurons (from Jerber et al, 2021), FDR < 5%

Both tables are uploaded as csv data files, with fields defined by the table below.

| | |
|---|---|
| gene_name | Hgnc (HUGO Gene Nomenclature Committee) symbol |
| ensembl_gene_id | Ensembl ID GRCh37 |
| pv_raw | Raw (nominal) p-value |
| pv | P-value corrected at gene level (Bonferroni) |
| qv | Q-value (globally corrected p-value using Storey procedure) |
| beta_G | Estimated conventional (persistent) effect size |
| beta_GxC | Estimated magnitude of allelic effects due to GxC |

### Supplementary Figures

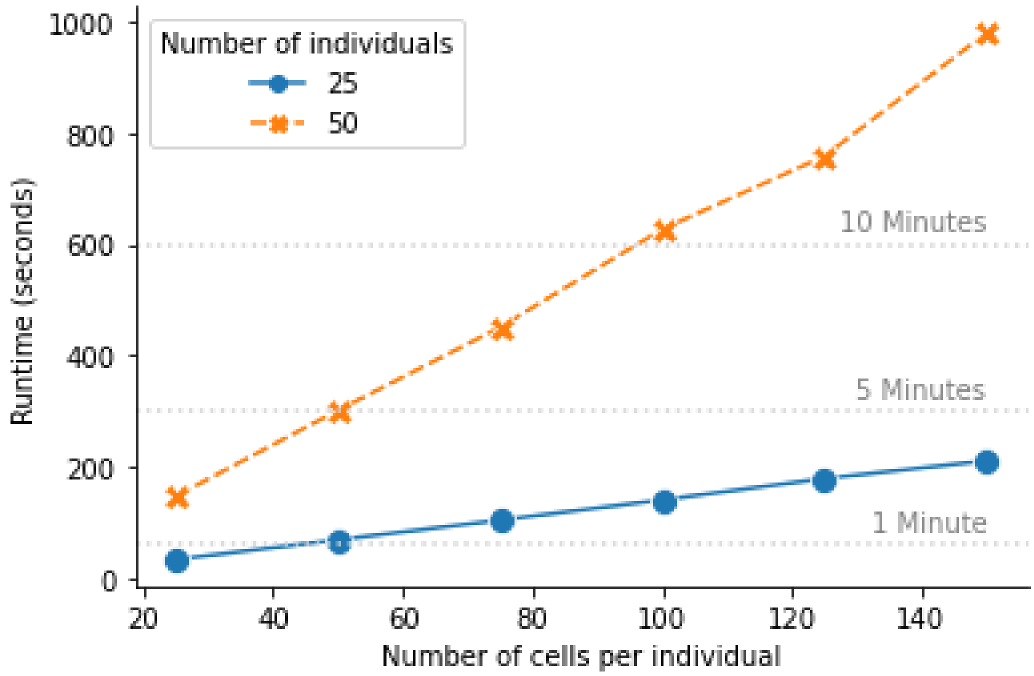


**Supplementary Figure 1.1: Empirical assessment of the computational complexity of CellRegMap.** Shown are empirical runtimes for simulated data (1 gene and 25 tested SNPs) for increasing number of cells observed per individual. CellRegMap scales linearly with the number of cells per individual. Runtimes evaluated on a 2018 MacBook Pro with 2,3 GHz Quad-Core Intel Core i5 processor.

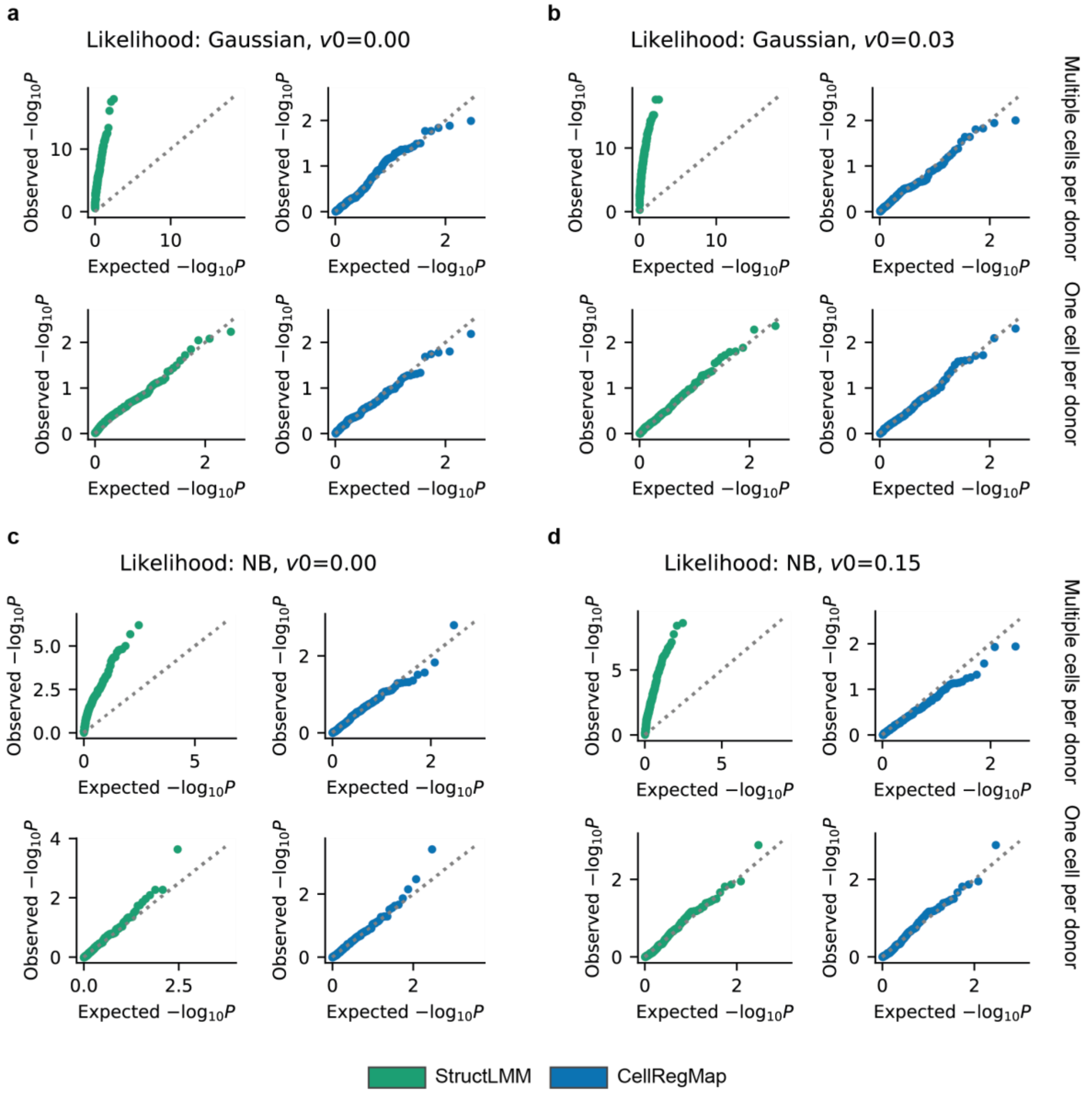


**Supplementary Figure 2.1: Assessment of statistical calibration on simulated data. (a-d)** QQ plots displaying expected versus observed negative log *P*-values for StructLMM and CellRegMap for a single gene and 200 tested SNPs. v0 denotes the fraction of variance explained by a persistent (non GxC) genetic effect for each SNP. Top rows of each panel show calibration when repeat structure is present (50 cells sampled in each of 50 individuals), while the bottom rows show results for no repeat structure (2,500 individuals, 1 cell sampled in each individual). **(a)** Simulated Gaussian residual noise, no persistent genetic effect. **(b)** Simulated Gaussian residual noise, with persistent genetic effect. **(c)** Simulated negative binomial noise, no persistent genetic effect. **(d)** Simulated negative binomial noise, with persistent genetic effect.

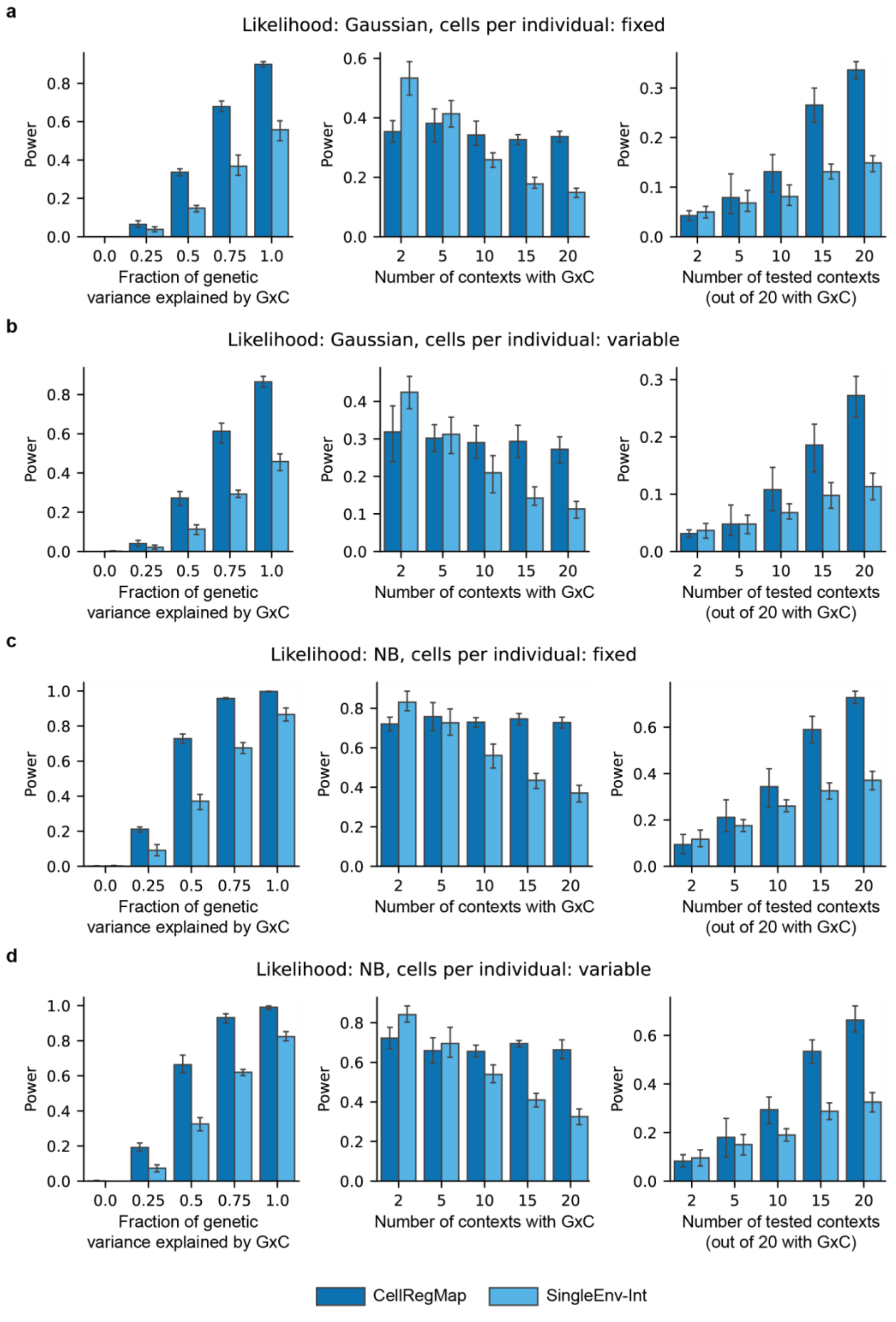


**Supplementary Figure 2.2: Additional results from the power assessment of CellRegMap on simulated data. (a-d)** Power as a function of the fraction of genetic variance explained by GxC, the number of simulated contexts with GxC and the number of tested cellular environments (analogous to main text Figure 2c-e), when simulating a fixed number of cells per donor (**a**, as in main text Fig. 2), as well as variable number of cells per individual (**b**). The equivalent settings are repeated for simulating negative binomial distributed residual noise (**c,d; Methods**). Error bars correspond to standard deviations estimated from 10 random initializations of the simulation procedure.

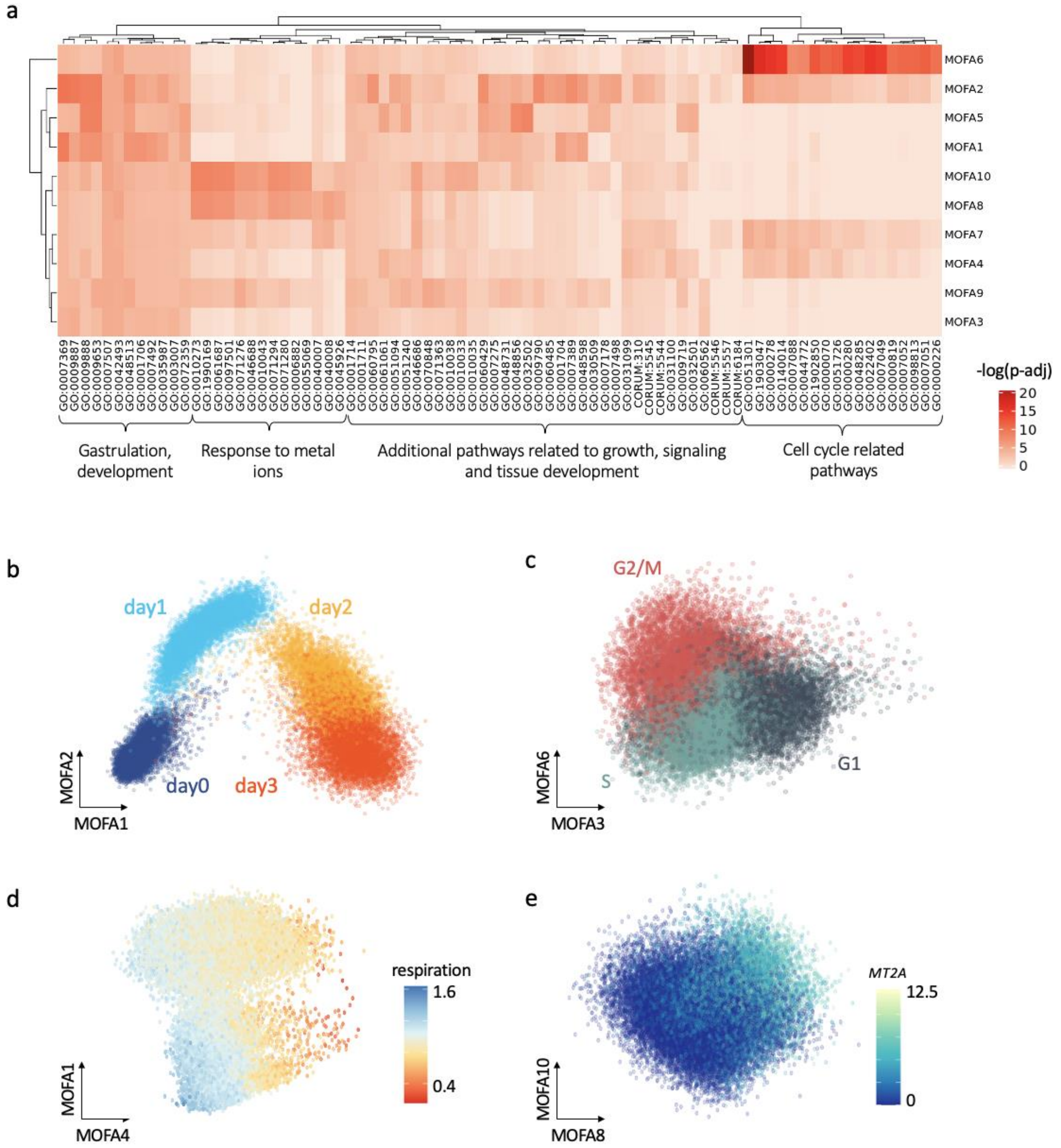


**Supplementary Figure 3.1: Annotation of the MOFA factors from the endoderm differentiation study.** **(a)** Heatmap displaying negative log p-values from GO enrichment analyses based on the absolute values of the of loadings of individual MOFA factors (Methods). **(b)** Scatter plot of MOFA factors 1 & 2, with colour corresponding to the time point of collection (day 0,1,2 & 3 of endoderm differentiation). **(c)** Scatter plot of MOFA factors 3 & 6, coloured by estimated cell cycle phase (G1, G2/M, S; estimated using Seurat; Methods). **(d)** Scatter plot of MOFA factors 1 & 4 capturing respiration (Methods). **(e)** Scatter plot of MOFA factors 8 & 10, capturing a signature linked to response to metal ions, coloured by expression of gene with top loadings, MT2A (Methods).

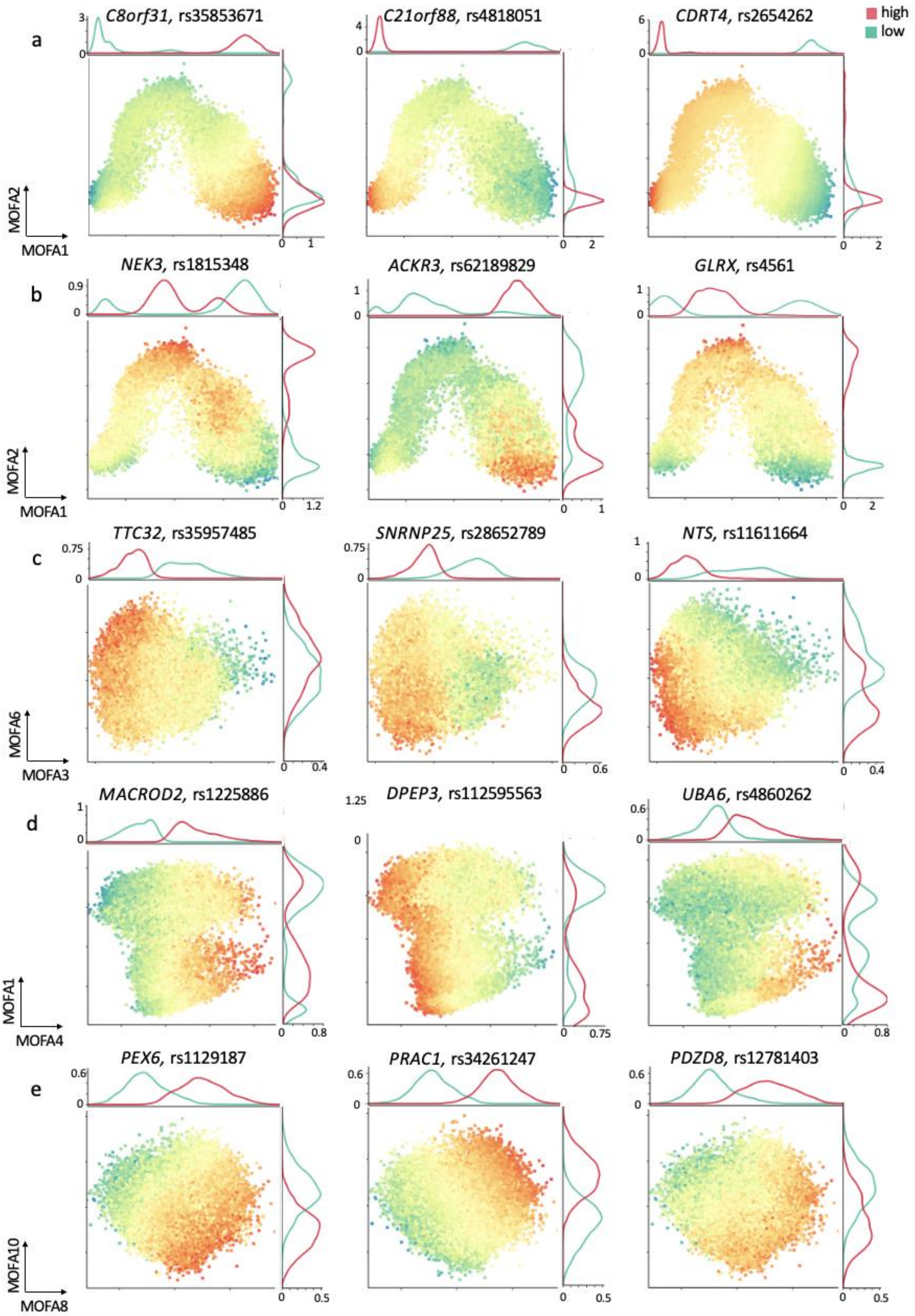


**Supplementary Figure 3.2: Additional examples of GxC interactions identified in the endoderm differentiation study using CellRegMap. (a)** Analogous to main text Fig. 3b (middle), examples of dynamic eQTL that vary continuously along MOFA 1 only. **(b)** Similar to main text Fig. 3b (right), additional examples of eQTL with GxC effects associated with MOFA factors 1 & 2. **(c)** Additional examples as in main text to Fig. 3c, displaying GxC effects across MOFA factors 3 & 6 (capturing cell cycle). **(d)** Additional examples as in main text Fig. 3c, displaying GxC effects along MOFA factors 4 and 1 (capturing cell respiration). **(e)** Examples of eQTL with GxC effects that are associated with cell states captured by MOFA 8 & 10, related to response to metal ions (c.f. Fig. 3.1a).

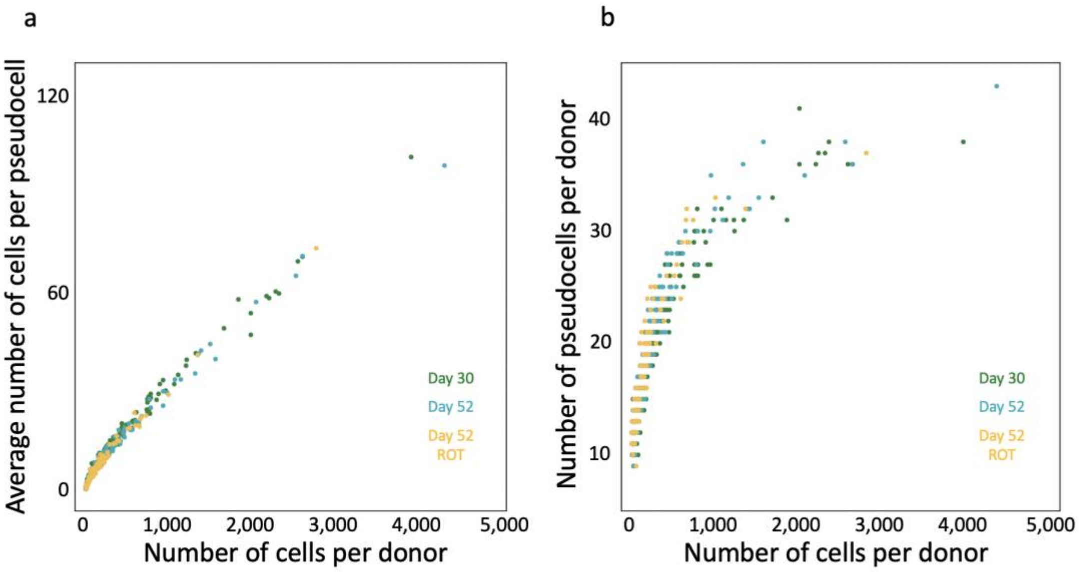


**Supplementary Figure 4.1. Dopaminergic neuron pseudocell aggregation. T**ranscriptionally related cells were aggregated into pseudocells, thereby reducing the sparsity in this dataset. Pseudocells were constructed separately for each individual and conditions (day 30, day 52, rotenone-treated day 52, Methods). **(a)** Scatter plot between the number of cells for each individual (x axis) versus the average number of cells contained in a pseudocell for the corresponding individual (y axis). Colour denotes the three main cell populations (collected across three conditions). **(b)** Analogous as in **a**, comparing the number of cells per donor versus the number of pseudocells per donor.

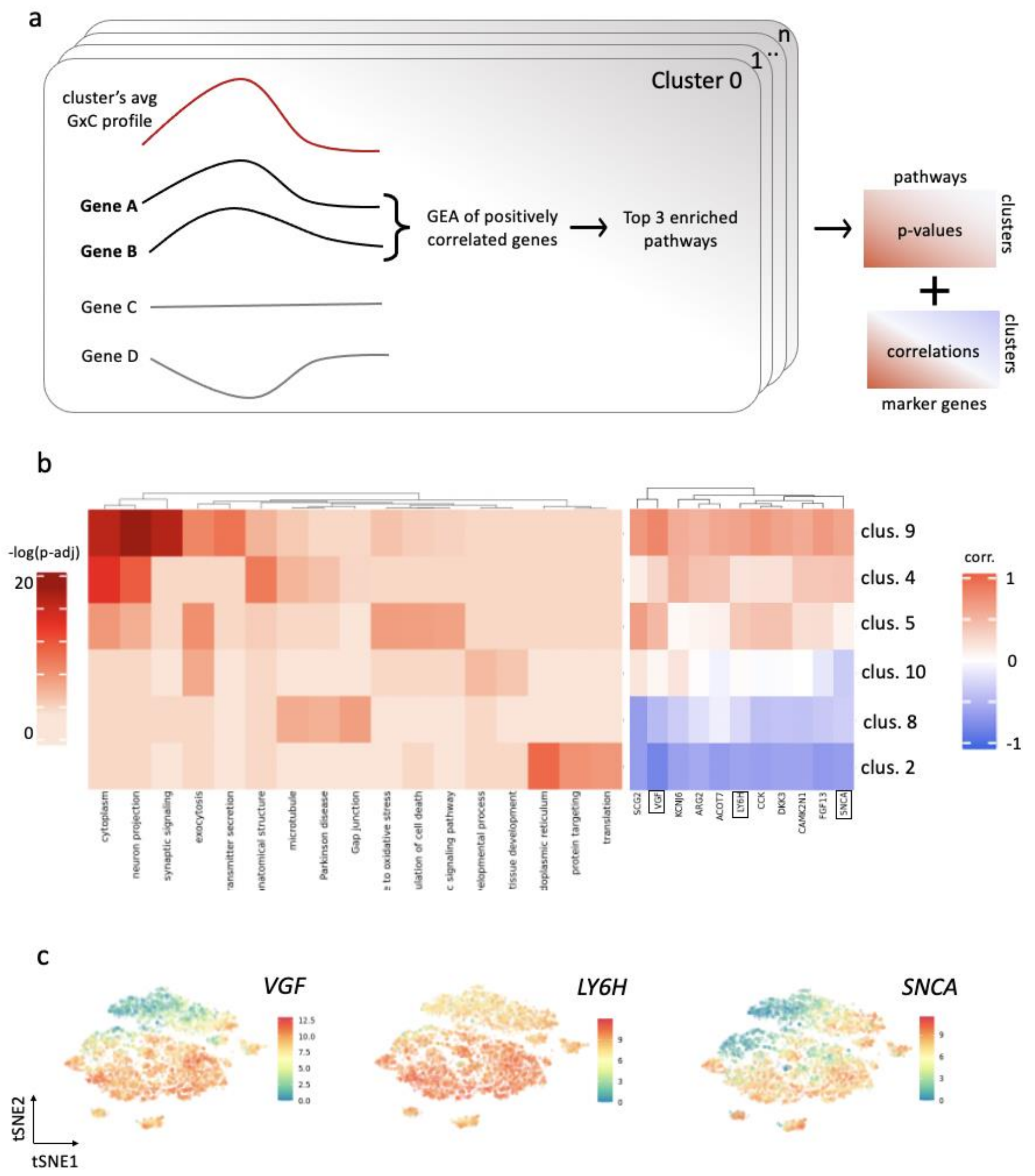


**Supplementary Figure 4.2. Annotation of GxC clusters based on gene set enrichment. (a)** Illustration of the clustering approach to identify principal patterns of GxC interactions. Briefly, for each cluster we considered genes whose single-cell profiles were positively correlated with the average GxC allelic effect profile. Genes with positive correlation (Pearson's correlation > 0.4) were considered for gene enrichment analysis using gprofiler, and up to 3 top significant (adjusted p-values < 0.05) terms were considered per cluster (Methods). **(b)** Left: Enrichment results for the 6 clusters in Fig. 4c-h. Shown is a heatmap of adjusted negative log p-values of enrichment results obtained by gprofiler. Right: Correlation coefficients between the GxC allelic effect profile and expression level for selected literature-curated dopaminergic neuron markers. For each cluster, the correlation coefficient between the expression level of the respective marker gene and the aggregate cluster allelic effect size profile is shown. **(c)** Expression profiles across pseudocells for three selected neuronal marker genes highlighted in the right panel of **b.** tSNE plots, coloured by expression level of the three genes.

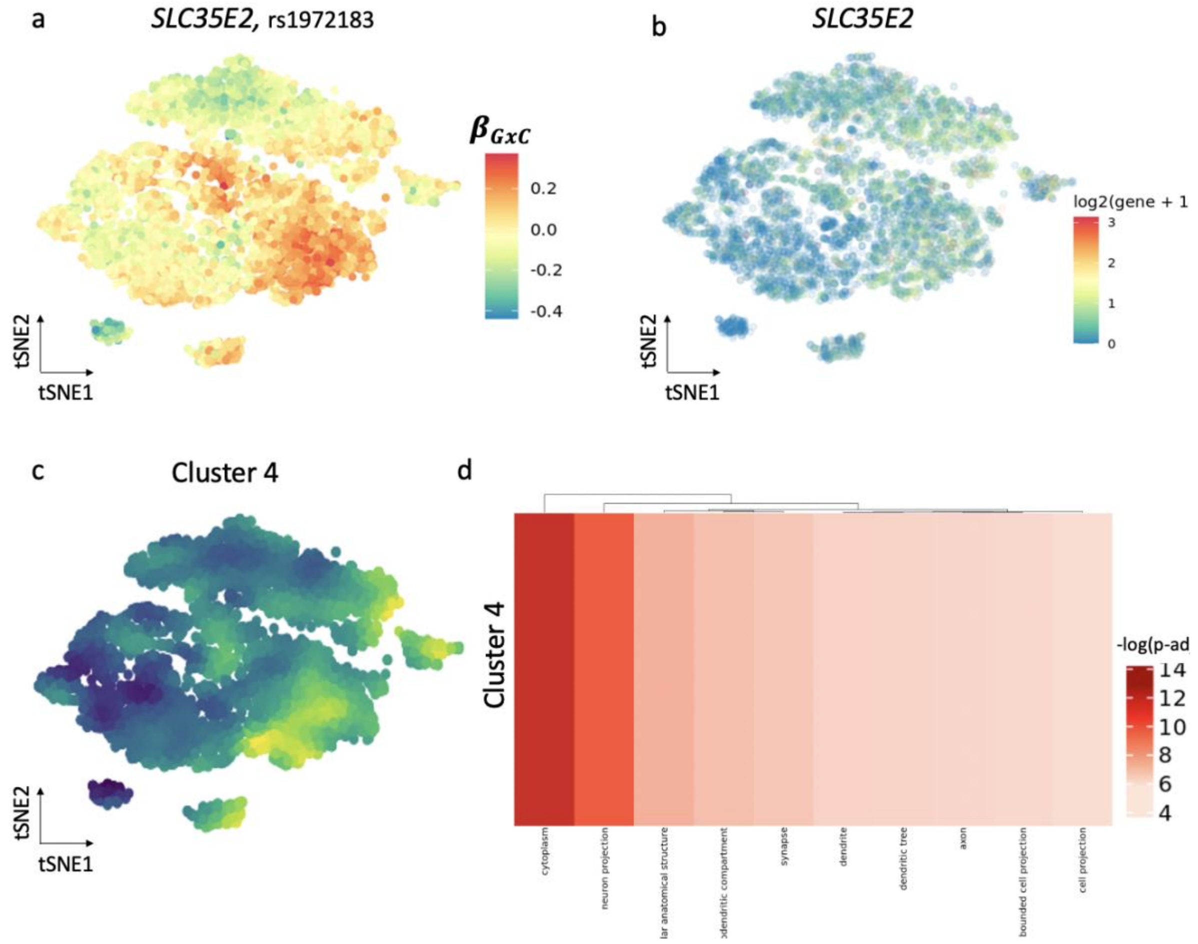


**Supplementary Figure 5.1. CellRegMap to fine-map cellular context for human disease variant.** **(a)** GxC profile at rs1972183 for *SLC35E2*, which is colocalized with a GWAS variant for sleeplessness and insomnia in the subpopulation of day 52 untreated cells. Shown is a scatter plot of the first two tSNE coordinates with colour denoting the estimated GxC effect component $\beta_{GxC}$. **(b)** As in **a**, with colour denoting *SLC35E2* single-cell gene expression levels. **(c)** Manifold of consensus relative GxC effect sizes estimates for cluster 4, which contains the GxC effects for *SLC35E2*. **(d)** Extract of the gene enrichment analysis for cluster 4 (based on **Supplementary Fig. 4.2b**).

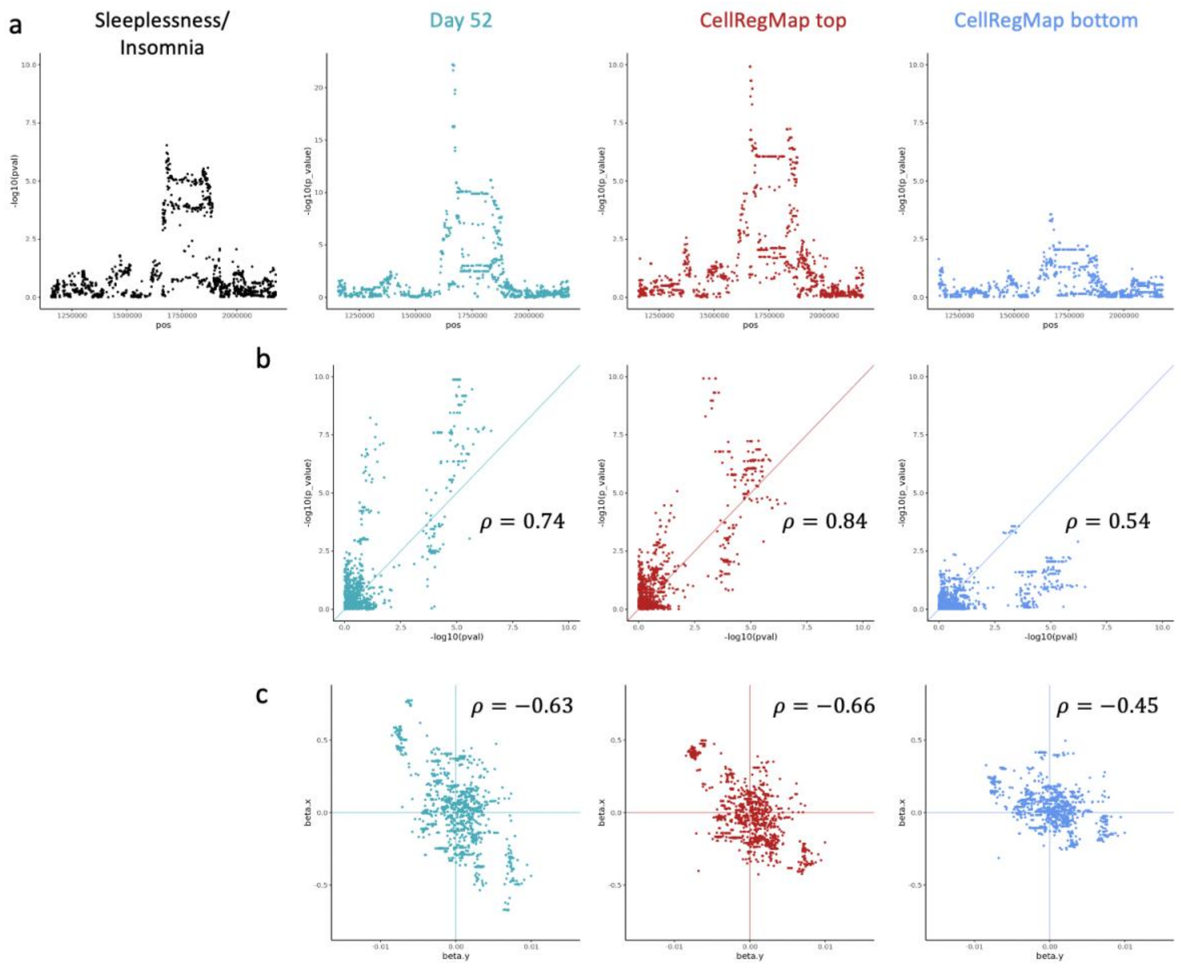


**Supplementary Figure 5.2. CellRegMap to fine-map a coloc variant.** GxC profile at rs1972183 for *SLC35E2* identifies cellular population linked to GWAS variant for insomnia/sleplessness. Columns represent all cells at day 52 (aqua), cell population that CellRegMap identifies to have strongest effects (red, top 30% quantile from $\beta_{GxC}$) and weakest (blue, bottom 30% quantile). **(a)** Manhattan plots. From left to right: original GWAS in the relevant genomic area, eQTL using aggregate expression estimates across all day 52 (untreated), eQTL Manhattan plot when considering day 52 cells in the top quantile, eQTL Manhattan plot when considering day 52 cells in the bottom quantile. **(b,c)** Comparison between GWAS signal and eQTL signal, considering alternative traits based on the cell populations as in a. **(b)** Scatter plots of negative log p-values from GWAS (x-axis) versus eQTL (y-axis). **(c)** As in b, displaying effect size estimates.
