## Supplementary Methods for "CellRegMap: A statistical framework for mapping context-specific regulatory variants using scRNA-seq"

### CellRegMap: A statistical framework for mapping context-specific regulatory variants using single cell RNA-sequencing.

#### Supplementary Methods

Anna S.E. Cuomo et al.

**8 Software availability** **26**

### 1 The CellRegMap model

CellRegMap builds on and extends the structured linear mixed model (StructLMM [1]), which has recently been proposed to test for genotype-environment interactions on physiological traits in population cohorts. CellRegMap extends this model to test for interactions between genotype and cellular context on gene expression using single-cell RNA-seq as readout. The model is not designed for variant discovery but instead designed to identify and characterize genotype-context (G×C) interactions at known expression quantitative trait loci (eQTL). In Section 1.1, the CellRegMap model is motivated and derived, followed by the introduction of an efficient scheme for parameter inference and statistical testing.

#### 1.1 Model definition

Commonly used methods for eQTL mapping based on linear or linear mixed models (LMM) relate individual genetic variants to the expression level of a gene of interest, i.e.,

$$\mathbf{y} = \mathbf{W}\boldsymbol{\alpha} + \mathbf{g}\beta_G + \boldsymbol{\psi}. \tag{1}$$

Here, $\mathbf{y}$ denotes a $N \times 1$ vector of the expression level of a gene of interest across $N$ individuals (typically measured using bulk RNA sequencing); $\mathbf{W}$ is the $N \times P$ design matrix of covariates and $\alpha_i, i = 1, .., P$ are the corresponding weights (note that in the main text, i.e., Figure 1d, covariates are omitted for brevity); $\mathbf{g}$ is the $N \times 1$ vector of alleles for each individual at the locus to be assessed and $\beta_G$ denotes the corresponding effect size. Finally $\boldsymbol{\psi}$ denotes i.i.d. noise, $\boldsymbol{\psi} \sim \mathcal{N}(\mathbf{0}, \sigma_n^2\mathbf{I})$. Additional random effect components have been introduced to account for relatedness between individuals [2], or to adjust for additive confounding sources of variation on gene expression [3].

###### Review of StructLMM

StructLMM [1] extends the conventional linear association test in Eq. (1) by including an additional random effect component to account for heterogeneity in effect sizes across individuals due to context-specific genetic effects. Briefly, StructLMM can be cast as:

$$\mathbf{y} = \mathbf{W}\boldsymbol{\alpha} + \mathbf{g}\beta_G + \mathbf{g} \odot \boldsymbol{\beta_{GxC}} + \mathbf{c} + \boldsymbol{\psi}, \tag{2}$$

where $\mathbf{g} \odot \boldsymbol{\beta_{GxC}}$ accounts for GxE and $\mathbf{c}$ for additive contributions of environmental variation. Unlike in conventional interaction tests, genotype-context interactions due to environmental variation are accounted by introducing an additional genetic effect with per-individual effects, where each individual has its own effect size. The symbol $\odot$ denotes the Hadamard product and $\boldsymbol{\beta_{GxC}} = [\beta_{GxC_1}, .., \beta_{GxC_N}]^T$ is a vector of per-individual effect sizes to account for heterogeneous genetic effects. Instead of explicitly estimating the GxC effect sizes, these parameters are marginalised under a multivariate normal prior distribution that is defined by an environmental context covariance matrix

$$\boldsymbol{\beta_{GxC}} \sim \mathcal{N}(\mathbf{0}, \sigma^2_{GxC}\mathbf{\Sigma}). \tag{3}$$

In this case the covariance $\mathbf{\Sigma}$ does not encode relatedness as in a classical LMM, but instead accounts for sample covariance due to different environmental states. The same environmental covariance matrix can also be used to parameterize a second random effect component to account for additive environmental contributions,

$$\mathbf{c} \sim \mathcal{N}(\mathbf{0}, \sigma^2_c\mathbf{\Sigma}). \tag{4}$$

Notably, StructLMM does not account for any additional structure across samples, such as population structure or repeat measurements. While discrete populations can be accounted for as covariates, more subtle relatedness or repeat structure cannot be effectively encoded using fixed effect covariates. Consequently, the model is not suitable for any experimental design where multiple observations are available for the same individual, which is the case in single-cell genetic studies where multiple cells are assayed from each individual. Such a repeat structure results in un-calibrated p-values; see also the results from the simulation study to assess calibration (main text Fig. 2).

**CellRegMap**

CellRegMap extends StructLMM to allow for applications to single-cell expression data by introducing an additional random effect component that accounts for relatedness or sample repeat structure. First we note that the phenotype $\mathbf{y}$ now represents single-cell resolved expression data, so our samples are expression levels in cells, not individuals. This introduces additional structure in the data, as typically multiple cells are sampled from the same individual. To account for this structure, an additional random effect component $\mathbf{u}$ is included in the model:

$$\mathbf{y} = \mathbf{W}\boldsymbol{\alpha} + \mathbf{g}\beta_G + \mathbf{g} \odot \boldsymbol{\beta_{GxC}} + \mathbf{c} + \mathbf{u} + \boldsymbol{\psi}. \tag{5}$$

Here, $\mathbf{y}$ has the cardinality of single-cells rather than individuals, and hence the G$\times$C component ($\mathbf{g} \odot \boldsymbol{\beta_{GxC}}$) accounts for interactions with cellular states and contexts (which are well defined at the level of single cells) as well as environmental exposures and stimuli (which can also be individual-level). The terms $\mathbf{c} \sim \mathcal{N}(\mathbf{0}, \sigma^2_c\mathbf{\Sigma})$ and $\boldsymbol{\psi} \sim \mathcal{N}(\mathbf{0}, \sigma^2_n\mathbf{I})$ have the same meaning as previously, noting that for these $N \times 1$ vectors $N$ now represents the total number of cells, not the number of unique individuals. Similarly, $\mathbf{\Sigma}$) is $N \times N$ and is again defined in the space of cells. The symbol $\boldsymbol{\beta_{GxC}} \sim \mathcal{N}(\mathbf{0}, \sigma^2_{GxC}\mathbf{\Sigma})$ now represents cell-level effect sizes, which again captures variation in genetic effects across cells. The additional random effect component $\mathbf{u}$ accounts for relatedness or the repeat structure, which is parameterized as a product kernel between relatedness ($\mathbf{R}$) and the environmental covariance ($\boldsymbol{\Sigma}$):

$$\mathbf{u} \sim \mathcal{N}(\mathbf{0}, \sigma_{rc}^2 \mathbf{R} \odot \boldsymbol{\Sigma}). \tag{6}$$

Here, $\mathbf{R}$ denotes the relatedness matrix of individuals expanded to all cells based on the known assignment of cells to individuals and the covariance $\boldsymbol{\Sigma}$ again denotes the cell-level environmental context. Notably, this parametrization extends the classical LMM, which would exclusively consider a relatedness component $\mathbf{R}$. One way to interpret this covariance is to account for polygenic interactions between environmental and relatedness, which has previously been considered to estimate the GxE component of heritability [4].

#### 1.2 Construction of the cellular context covariance

Typically, we define $\boldsymbol{\Sigma} = \mathbf{E}\mathbf{E}^T$, and hence the cellular context covariance is a linear function of a matrix of environmental states $\mathbf{E}$. In practice, we consider as cellular contexts axes of variation in the dataset (for example captured by principal components or MOFA [5] factors), appropriately standardized (mean=0, standard deviation=1) and build $\boldsymbol{\Sigma} = \mathbf{E}\mathbf{E}^T$ accordingly. Depending on the type and structure of cellular contexts, $\boldsymbol{\Sigma}$ can simply separate cells into groups, and appear as a block diagonal or capture continuous transitions (main text Fig. 1c). In principle, CellRegMap can also be use in conjunction with other parameterizations of the cell context covariance.

#### 1.3 Statistical testing

The main operations in the CellRegMap model, including parameter inference and tests are implemented in its marginalised form. Integrating over the random effect components, the marginal likelihood of the model in Eq. (5) follows as:

$$\mathbf{y} \sim \mathcal{N}(\mathbf{W}\boldsymbol{\alpha} + \mathbf{g}\beta_G, \sigma_{GxC}^2 \text{diag}(\mathbf{g})\boldsymbol{\Sigma}\text{diag}(\mathbf{g}) + \sigma_c^2 \boldsymbol{\Sigma} + \sigma_{rc}^2 \mathbf{R} \odot \boldsymbol{\Sigma} + \sigma_n^2 \mathbf{I}). \tag{7}$$

While in principle CellRegMap can be used to test for different components, including additive genetics, interactions or the combination of both effects (see [1] for details on StructLMM), we here focus on GxC effects. In order to evaluate the significant contribution of GxC effects, we consider a statistical test that compares the following hypotheses, from Eq.(5):

$$H_0 : \sigma_{GxC}^2 = 0,$$

$$H_1 : \sigma_{GxC}^2 > 0.$$

We use Rao's Score test [6] to evaluate significance, which allows us to only calculate the MLE[1] of the parameters under the null hypothesis $H_0$, which follows from Eq.(7) as:

$$\mathbf{y}|H_0 \sim \mathcal{N}(\mathbf{W}\boldsymbol{\alpha} + \mathbf{g}\beta_G, \sigma_c^2\boldsymbol{\Sigma} + \sigma_{rc}^2\mathbf{R} \odot \boldsymbol{\Sigma} + \sigma_n^2\mathbf{I}). \tag{8}$$

To test for GxC interactions we adapt the score-based testing scheme employed in StructLMM, which in turn adopts fast LMM testing in LMMs as first proposed in Lippert et al., [7]. We here review the key steps involved.

First, we define the score-based test statistic Q as:

$$\mathrm{Q} = \frac{1}{2}\mathbf{y}^T\mathbf{P}_0\frac{\partial\mathbf{K}}{\partial\theta}\mathbf{P}_0\mathbf{y}, \tag{9}$$

where $\mathbf{K}$ denotes the full covariance matrix, i.e., from Eq. (5):

$$\mathbf{K} = \sigma_{GxC}^2\mathrm{diag}(\mathbf{g})\boldsymbol{\Sigma}\ \mathrm{diag}(\mathbf{g}) + \sigma_c^2\boldsymbol{\Sigma} + \sigma_{rc}^2\mathbf{R} \odot \boldsymbol{\Sigma} + \sigma_n^2\mathbf{I}, \tag{10}$$

and

$$\mathbf{P}_0 = \mathbf{K}_0^{-1} - \mathbf{K}_0^{-1}\mathbf{X}(\mathbf{X}^T\mathbf{K}_0^{-1}\mathbf{X})^{-1}\mathbf{X}^T\mathbf{K}_0^{-1} \tag{11}$$

is a matrix that projects out the fixed effects [7, 8]. In our case (Eq.(5)), the fixed effects include covariates and the persistent effect of the variant tested: $\mathbf{X} = [\mathbf{W}, \mathbf{g}]$, and $\mathbf{K}_0$ is as in Eq.(16). Using Eq.(10) and considering the parameter $\theta = \sigma_{GxC}^2$, we can derive:

$$\frac{\partial\mathbf{K}}{\partial\sigma_{GxC}^2} = \mathrm{diag}(\mathbf{g})\boldsymbol{\Sigma}\ \mathrm{diag}(\mathbf{g}). \tag{12}$$

Next, let us define $\mathbf{K}_1 = \mathrm{diag}(\mathbf{g})\boldsymbol{\Sigma}\ \mathrm{diag}(\mathbf{g})$; substituting in Eq.(9), we can write:

$$\mathrm{Q} = \frac{1}{2}\mathbf{y}^T\mathbf{P}_0\mathbf{K}_1\mathbf{P}_0\mathbf{y}. \tag{13}$$

As before, $H_1 : \sigma_{GxC}^2 > 0$, noting that as a variance parameter, $\sigma_{GxC}^2$ is constrained to take on positive values. As a result, the score test statistic Q follows a mixture of $\chi^2$ distributions[2]:

$$\mathrm{Q} \sim \sum_i \lambda_i\chi_1^2, \tag{14}$$

[1] maximum likelihood estimator

[2] We refer the reader specifically to the supplementary methods from [8] for a proof.

where $\lambda_i$'s are the non-zero eigenvalues of $\frac{1}{2}\mathbf{P}_0^{\frac{T}{2}}\frac{\partial \mathbf{K}}{\partial \theta}\mathbf{P}_0^{\frac{1}{2}}$.

It can be shown that for a matrix $\mathbf{A}$ the non-zero eigenvalues of $\mathbf{A}\mathbf{A}^T$ are the same as those of $\mathbf{A}^T\mathbf{A}$ , thus we can re-arrange and compute $\lambda_i$'s as the eigenvalues of:

$$\frac{1}{2}\frac{\partial \mathbf{K}}{\partial \theta}^{\frac{T}{2}}\mathbf{P}_0\frac{\partial \mathbf{K}}{\partial \theta}^{\frac{1}{2}} \tag{15}$$

instead. To evaluate the significance of the score-best test statistic Q we use the approach described in Sequence Kernel Association Test (SKAT [9]), thereby using the Davies exact method [10] to compute the corresponding p values, and switching to the modified moment matching approximation method [11, 12, 13] when this fails to converge.

##### 1.4 Implementation

To enable efficient parameter inference, we extended the strategy in StructLMM [1], which builds on the reparametrization of the LMM likelihood proposed in [7]. Briefly, the key is to rewrite the overall covariance of $\mathbf{y}|H_0$ in the form: $\sigma_m^2(\mathbf{M}+\delta\mathbf{I})$, where $\mathbf{M}$ is ideally low rank to enable an efficient singular value decomposition. To do so, we introduce a weight parameter $\rho_1$, such that the covariance matrix of $\mathbf{y}$ under the null hypothesis, $\mathbf{K}_0 = \text{Cov}(\mathbf{y}|H_0)$ can be cast as:

$$\begin{aligned}\mathbf{K}_0 = \sigma_c^2\mathbf{\Sigma} + \sigma_{rc}^2\mathbf{R}\odot\mathbf{\Sigma} + \sigma_n^2\mathbf{I} = \\ \sigma_m^2\ [\rho_1\mathbf{\Sigma} + (1-\rho_1)\mathbf{R}\odot\mathbf{\Sigma}] + \sigma_n^2\mathbf{I} = \\ \sigma_m^2\{\mathbf{M}(\rho_1) + \delta_1\mathbf{I}\},\end{aligned} \tag{16}$$

where $\sigma_m^2\rho_1 = \sigma_c^2$, $\sigma_m^2(1-\rho_1) = \sigma_{rc}^2$, $\delta_1 = \sigma_n^2/\sigma_m^2$, and $\mathbf{M}(\rho_1) = \rho_1\mathbf{\Sigma} + (1-\rho_1)\mathbf{R}\odot\mathbf{\Sigma}$.

We note that $\mathbf{M}$ only depends on $\rho_1$. The parameter $\rho_1$ is optimized using a grid search with values in the range 0 to 1, selecting the value of $\rho_1$ that maximises the likelihood of $\mathbf{y}|H_0$ under the null. Once $\rho_1$ is fixed, the decomposition of $\rho_1\mathbf{\Sigma} + (1-\rho_1)\mathbf{R}\odot\mathbf{\Sigma}$ can be found using the observation that the decomposition of the linear combination of two square symmetric matrices, e.g.,:

$$\mathbf{M} = a\mathbf{A} + b\mathbf{B} \tag{17}$$

can be found as a function of the decomposition of the two original matrices, i.e.,:

$$\mathbf{N} = [\sqrt{a}\mathbf{C}|\sqrt{b}\mathbf{D}] \tag{18}$$

such that $\mathbf{M} = \mathbf{N}\mathbf{N}^T, \mathbf{A} = \mathbf{C}\mathbf{C}^T$ and $\mathbf{B} = \mathbf{D}\mathbf{D}^T$.

In our case, $\mathbf{A} = \mathbf{\Sigma}$ and $\mathbf{B} = \mathbf{R} \odot \mathbf{\Sigma}$. If $\mathbf{\Sigma} = \mathbf{E}\mathbf{E}^T$, its decomposition is straight-forward, i.e., $\mathbf{C} = \mathbf{E}$. The decomposition of the second term $(\mathbf{R} \odot \mathbf{\Sigma})$, assuming $\mathbf{\Sigma} = \mathbf{E}\mathbf{E}^T$ and $\mathbf{R} = \mathbf{G}\mathbf{G}^T$ is a bit less trivial. Assuming that the rank of $\mathbf{R}$ is the number of individuals, that the rank of $\mathbf{\Sigma}$ is the number of environments, and that the latter is smaller than the former, we can obtain $\mathbf{D}$ as the decomposition of $\mathbf{R} \odot \mathbf{\Sigma}$ by:

$$\mathbf{R} \odot \mathbf{\Sigma} = \mathbf{R} \odot \sum_i [\mathbf{v}_i \lambda_i \mathbf{v}_i^T] = \sum_i [\mathbf{R} \odot (\mathbf{v}_i \lambda_i \mathbf{v}_i^T)]. \tag{19}$$

The terms in the sum can be rewritten as:

$$\begin{aligned}
\mathbf{R} \odot (\mathbf{v}_i \lambda_i \mathbf{v}_i^T) &= \mathbf{R} \odot (\mathbf{u}_i \mathbf{u}_i^T) \\
&= \mathbf{R} \odot (\text{diag}(\mathbf{u}_i)\mathbf{e}\mathbf{e}^T \text{diag}(\mathbf{u}_i)) \\
&= \text{diag}(\mathbf{u}_i)[\mathbf{R} \odot \mathbf{e}\mathbf{e}^T]\text{diag}(\mathbf{u}_i) \\
&= \text{diag}(\mathbf{u}_i)\mathbf{R}\text{diag}(\mathbf{u}_i) \\
&= \text{diag}(\mathbf{u}_i)\mathbf{G}\mathbf{G}^T\text{diag}(\mathbf{u}_i) \\
&= \mathbf{Lk}_i \mathbf{Lk}_i^T,
\end{aligned} \tag{20}$$

where $\mathbf{Lk}_i = \text{diag}(\mathbf{u}_i)\mathbf{G}$; $\lambda_i$'s and $\mathbf{v}_i$'s are eigenvalues and eigenvectors of $\mathbf{E}\mathbf{E}^T$; $\mathbf{u}_i = \sqrt{\lambda_i}\mathbf{v}_i$ and $\mathbf{e}$ is a vector of ones ($\mathbf{e} = [1..1]$).

Thus, $\mathbf{M} = \mathbf{\Sigma} + \mathbf{R} \odot \mathbf{\Sigma}$ can be written as the following sum:

$$\mathbf{M} = \mathbf{\Sigma} + \sum_i \mathbf{Lk}_i \mathbf{Lk}_i^T, \tag{21}$$

from which follows:

$$\mathbf{N} = [\mathbf{E} \; |\mathbf{Lk}_1 \; |.. \; |\mathbf{Lk}_{ne}], \tag{22}$$

where $ne$ is the number of contexts considered (and the rank of $\mathbf{\Sigma}$).

#### 1.5 Computational complexity

From using the implementation described above, it follows that the complexity is O(N) where N is the minimum between the number of cells and the product of number of unique individuals $\times$ the number of cellular contexts. Runtimes were also evaluated empirically, using simulated phenotypes for one gene, 25 SNPs and 20 different contexts (see section 3). We considered 25 and 50 individuals, respectively, and assessed computational runtimes as a function of the number of cells per individual (25, 50, 75, 100, 125, 150) using a 2018 MacBook Pro with 2,3 GHz Quad-Core Intel Core i5 processor (Supp. Fig. 1.1).

#### 2 Predicting cell-specific effect sizes driven by GxC interactions

Using CellRegMap it is possible to estimate cell-lelel allelic effects due to GxC, (thus estimating $\boldsymbol{\beta_{GxC}}$ from eq. (7)) for each gene-SNP pair tested. For this derivation, we will use the function representation of linear mixed model as a Gaussian process ($\mathcal{GP}$). Briefly (for more details see section 7.1),

$$\mathbf{f} \sim \mathcal{GP}\{\mathbf{m}; k(\mathbf{X}, \mathbf{X})_{\boldsymbol{\theta}}\}, \tag{23}$$

or:

$$\mathbf{f} = \mathbf{m} + \mathbf{c}, \tag{24}$$

where we define:

$$\mathbf{m} = \mathbf{X}\boldsymbol{\beta}_{\theta} \text{ and } \mathbf{c} \sim \mathcal{N}(\mathbf{0}, k(\mathbf{X}, \mathbf{X})_{\theta}). \tag{25}$$

When we model our data ($\mathbf{X}$ and $\mathbf{y}$), we consider $\mathbf{y}$ as being a sample from the random variable $\mathbf{f}$, and $\mathbf{X}$ as being fixed. Let us collectively call the parameters $\boldsymbol{\theta}$ and $\hat{\boldsymbol{\theta}}_{BLUP}$ (or $\hat{\boldsymbol{\theta}}$) their best linear unbiased predictor (BLUP) estimator [14].

For out of sample ($*$) prediction, we can define [15]:

$$\mathbf{y}_* \coloneqq \mathrm{E}[\mathbf{f}_*|\mathbf{y}]_{\hat{\boldsymbol{\theta}}}, \tag{26}$$

which can be written as (see section 7.1 for intermediate steps):

$$\mathbf{y}_* = \mathbf{m}_* + k(\mathbf{X}_*, \mathbf{X})k(\mathbf{X}, \mathbf{X})^{-1}(\mathbf{y} - \mathbf{m}). \tag{27}$$

In our case, we use a modified model from eq. (5) where the additive environments are modelled as fixed effects:

$$\mathbf{y} = \overbrace{\mathbf{W}\boldsymbol{\alpha} + \mathbf{g}\beta_G + \mathbf{E}\boldsymbol{\gamma}}^{\mathbf{m}} + \overbrace{\mathbf{g} \odot \boldsymbol{\beta}_{GxE} + \mathbf{u} + \boldsymbol{\psi}}^{\mathbf{c}}. \tag{28}$$

By substituting $\mathbf{m}$ and $\mathbf{c}$ into eq. (27) we obtain a formula for $\mathbf{y}_*$:

$$\mathbf{y}_* = \mathbf{W}_*\boldsymbol{\alpha} + \mathbf{g}_*\beta_G + \mathbf{E}_*\boldsymbol{\gamma} + k(\mathbf{X}, \mathbf{X}_*)k(\mathbf{X}, \mathbf{X})^{-1}(\mathbf{y} - \mathbf{W}\boldsymbol{\alpha} + \mathbf{g}\beta_G + \mathbf{E}\boldsymbol{\gamma}). \tag{29}$$

Now, to obtain an estimate for $\boldsymbol{\beta}_*$, we can evaluate eq. (29) when $\mathbf{g}_* = 0$ ($\mathbf{y}_*(\text{ref})$) and when $\mathbf{g}_* = 1$ ($\mathbf{y}_*(\text{alt})$), and consider the difference, scaled by a number based on the MAF[3] ($p$):

$$\boldsymbol{\beta}^*_{GxC} = \frac{1}{\sqrt{2p(1-p)}} \left(\mathbf{y}_*(\text{alt}) - \mathbf{y}_*(\text{ref})\right), \tag{30}$$

from which, skipping a few steps:

$$\boldsymbol{\beta}^*_{GxC} = \frac{1}{\sqrt{2p(1-p)}} \left\{\mathbf{1}\beta_G + \sigma^2_{GxC}\mathbf{E}_*(\mathbf{g} \odot \mathbf{E})^T\mathbf{K}^{-1}(\mathbf{y} - \mathbf{W}\boldsymbol{\alpha} - \mathbf{g}\beta_G - \mathbf{E}\boldsymbol{\gamma})\right\}, \tag{31}$$

where $\mathbf{K} = \text{Cov}(\mathbf{y}) = \sigma^2_{GxE}(\mathbf{g} \odot \mathbf{E})(\mathbf{g} \odot \mathbf{E})^T + \sigma^2_{re}\mathbf{R} \odot \mathbf{E}\mathbf{E}^T + \sigma^2_n\mathbf{I}$.

By setting $\mathbf{E}_* = \mathbf{E}$, we can perform in-sample estimation of allelic effects.

[3] minor allele frequency

### 3 Simulation strategy

We used simulation experiments to show calibration and power of CellRegMap. The code for building simulations generated from the model can be found at `https://github.com/annacuomo/CellRegMap_analyses/tree/main/simulations`.

#### 3.1 Genotype data

Given $S$ number of SNPs, we first generated $S$ random minor allele frequencies (MAFs) between 0.2 and 0.45. Given those, and a number of donors $N$, we generated $S$ independent $N \times 1$ $\mathbf{g}$ vectors as random vectors of 0, 1, 2 with probabilities given by the corresponding MAF ($p(2) = MAF^2$, $p(0) = (1-MAF)^2$, and $p(1) = 1 - p(0) - p(2)$). These represent the genotype vectors.

#### 3.2 Modelling number of cells per donor

We considered three settings when modelling the number of cells per donor. For both calibration and power simulations, we considered a fixed number of cells per donor, i.e., 50 cells per donor for 50 donors, resulting in a total of 2,500 samples (cells). For the calibration experiments, we additionally considered simulations without such repeat structure, simulating only 1 cell for 2,500 donors. Lastly, to evaluate the effects of non-uniform numbers of cells on discovery power, we also simulated a variable number of cells. More specifically, the number of cells for donor $i \in \{1, \ldots, 50\}$ was calculated as

$$\text{round}(2i \cdot 50/(50+1)), \tag{32}$$

resulting again in 2,500 cells in total. The genotype vectors ($\mathbf{g}$'s) are expanded appropriately to account for the repeat structure (such that all cells from an individual are assigned the same genotype value).

#### 3.3 Relatedness matrix

We defined $\mathbf{R}$ as the repeatedness matrix, by building a block-diagonal matrix, where each block represents an individual and its size depends on the number of cells for that donor.

#### 3.4 Cellular Contexts

The cell-cell context covariance matrix was defined as $\mathbf{\Sigma} = \mathbf{E}\mathbf{E}^T$, where $\mathbf{E}$ corresponded to a subset of principal component (PC) embeddings calculated from scRNA-seq of differentiating iPS cell lines [16]. Unless otherwise stated, we used 20 PCs for all experiments. When the number of cells per donor was larger than one, we sampled cells from the full embedding matrix such that all cells for one donor shared the same original cell line.

#### 3.5 Phenotype

Phenotype vectors were simulated under both Gaussian and negative binomial noise models. Briefly, in the Gaussian case we simulated from the full model (Eq.(5)), using only an intercept as covariate, and a single SNP ($\mathbf{g}_G$) having a persistent effect on $\mathbf{y}$ and another ($\mathbf{g}_{GxC}$) having GxC effects, such that:

$$\mathbf{y} = \mathbf{y}_0 + \underbrace{\mathbf{g}_G \beta_G}_{\mathbf{y}_G} + \underbrace{\mathbf{g}_{GxC} \odot \boldsymbol{\beta}_{\boldsymbol{GxC}}}_{\mathbf{y}_{GxC}} + \underbrace{\mathbf{c}}_{\mathbf{y}_C} + \underbrace{\mathbf{u}}_{\mathbf{y}_{RC}} + \underbrace{\boldsymbol{\psi}}_{\mathbf{y}_n}. \tag{33}$$

##### 3.5.1 Variance terms

We column-normalised all terms so that the total variance sums to 1 (see details below). Next, we set the variance explained by both genetic terms $\text{var}_G + \text{var}_{GxC} = \sigma_0^2$, and call the rest $v = 1 - \sigma_0^2$. Further, we regulated the amount of variance driven by GxC using an additional weighting factor $\rho_0$, such that: $\text{var}_G = (1 - \rho_0)\sigma_0^2$ and $\text{var}_{GxC} = \rho_0 \sigma_0^2$. For simplicity, the variances explained by the last three terms were set to the same value: $\sigma_C^2 = \sigma_{RC}^2 = \sigma_n^2 = v/3$ for Gaussian and $\sigma_C^2 = \sigma_{RC}^2 = v/2, \sigma_n^2 = 0$ in the case of a negative binomial noise model, respectively, such that:

$$\mathbf{y} = \mathbf{y}_0 + \underbrace{\underbrace{\mathbf{g}_G \beta_G}_{\text{var}_G} + \underbrace{\mathbf{g}_{GxC} \odot \boldsymbol{\beta}_{\boldsymbol{GxC}}}_{\text{var}_{GxC}}}_{\sigma_0^2} + \underbrace{\underbrace{\mathbf{e}}_{\sigma_C^2} + \underbrace{\mathbf{u}}_{\sigma_{RC}^2} + \underbrace{\boldsymbol{\psi}}_{\sigma_n^2}}_{v}. \tag{34}$$

##### 3.5.2 Persistent genetic effects

First, we define an $S \times 1$ effect size vector $\boldsymbol{\beta}$, which represent the effect size of each of the $S$ SNPs modelled. We model most SNPs to have no effect $\beta_i = 0$ except for one 'causal' SNP ($\mathbf{g}_G$), which we randomly set to either 1 or $-1$ (i.e., $\beta_G = \pm 1$). Next, we scale the vector to ensure that the variance explained by $\mathbf{y}_G$ is $\text{var}_G$:

$$\boldsymbol{\beta}_s = \frac{1}{\sqrt{\text{var}_G}} \boldsymbol{\beta}.$$

Finally, we compute the persistent genetic effect component of the phenotype vector by multiplying the genotype matrix $\mathbf{G}$ by the resulting vector scaled $\boldsymbol{\beta}_s$, i.e.,:

$$\mathbf{y}_G = \mathbf{G} \boldsymbol{\beta}_s. \tag{35}$$

###### 3.5.3 GxC effects

For every SNP $i$, let $\mathbf{y}_{G\times C,i} = \mathbf{g}_i \odot (\mathbf{E}\boldsymbol{\alpha_i})$ be its G×C effect, where $\mathbf{E}$ is the $N \times n_E$ cellular environment matrix and $\boldsymbol{\alpha}_i$ is a $n_E \times 1$ normally distributed random variable such that:

$$\boldsymbol{\alpha}_i \sim \mathcal{N}(0, \sigma^2_{G\times C,i}\mathbf{I}_{n_E}). \tag{36}$$

Similarly to before, to ensure that the variance explained is $\mathrm{var}_{GxC}$, we set $\sigma^2_{GxC,i} = \mathrm{var}_{GxC}$ if $\mathbf{g}_i$ is causal (assuming once again a single SNP being causal for G×C effects) and $\sigma^2_{GxC,i} = 0$ otherwise. The G×C component of the phenotype can then be obtained as:

$$\mathbf{y}_{GxC} = \sum_i \mathbf{y}_{GxC,i} = \sum_i \mathbf{g}_i \odot \mathbf{E}\boldsymbol{\alpha_i}. \tag{37}$$

###### 3.5.4 Additive random effects

Next, we simulate the last three random effect terms, noting that we simulate them all to explain the same variance.

First, the effect of cellular contexts directly:

$$\mathbf{y}_C = \mathbf{E}(\sigma_C\ \mathbf{n}), \quad \text{with } \mathbf{n} \sim \mathcal{N}(\mathbf{0}, \mathbf{I}). \tag{38}$$

Next, the background term to account for relatedness ($\mathbf{R} \odot \mathbf{E}\mathbf{E}^T$):

$$\mathbf{y}_{RC} = [\mathbf{L}\mathbf{k}_1 ... \mathbf{L}\mathbf{k}_{n_E}](\sigma_{RE}\ \mathbf{n}), \quad \text{with } \mathbf{n} \sim \mathcal{N}(\mathbf{0}, \mathbf{I}), \tag{39}$$

where $\mathbf{Lk}$'s are as in Eq. (20).

###### 3.5.5 Noise

For Gaussian noise we model the noise term as:

$$\mathbf{y}_n = \sigma_n\ \mathbf{n}, \quad \text{with } \mathbf{n} \sim \mathcal{N}(\mathbf{0}, \mathbf{I}). \tag{40}$$

Alternatively, we consider a negative binomial noise model, where we set $\boldsymbol{\psi} = \mathbf{y}_n = \mathbf{0}$ and map $\mathbf{y}$ to the mean of a negative binomial distribution using a log link function. The probability mass function of a negative binomial is commonly parameterized as

$$P(k; r, p) = \frac{\Gamma(k + r)}{k!\Gamma(r)} p^r (1 - p)^k \tag{41}$$

where $k + r$ is the number of trials and $r$ is the number of successes for a probability of success $p$. We parameterize the distribution in terms of its mean $\mu$ and dispersion parameter $\phi$, such that

$$r = \phi^{-1} \tag{42}$$

$$p = \frac{r}{r + \mu} \tag{43}$$

The dispersion parameter of the negative binomial was set to $\phi = 1.5$ for all simulations, and the intercept was set to $y_0 = 2.5$.

### 4 Model validation using simulated data

In order to validate the calibration of CellRegMap and to assess statistical power of the model, we considered simulated data generated as described in Section 3. For comparison, we also considered two alternative models as described in the next section.

#### 4.1 Alternative methods

We considered different comparison methods for calibration and power assessment of our model.

##### 4.1.1 StructLMM

First, we assessed calibration of our method in comparison to the original StructLMM model, thus comparing StructLMM:

$$\mathbf{y} = \mathbf{g}\beta_G + \mathbf{g} \odot \boldsymbol{\beta_{GxC}} + \mathbf{c} + \boldsymbol{\psi} \tag{44}$$

vs CellRegMap:

$$\mathbf{y} = \mathbf{g}\beta_G + \mathbf{g} \odot \boldsymbol{\beta_{GxC}} + \mathbf{c} + \mathbf{u} + \boldsymbol{\psi}, \tag{45}$$

where $\mathbf{y}$, $\mathbf{g}$, $\beta_G$, $\mathbf{g}$, $\boldsymbol{\beta_{GxC}}$, $\mathbf{c}$, $\mathbf{u}$ and $\boldsymbol{\psi}$ are as previously described (Eqs. (2), (5)). This comparison was conducted to confirm the need for the relatedness component (using a random effect component $\mathbf{u}$).

##### 4.1.2 SingleEnv-Int

For power analyses, we compared CellRegMap to one more traditionally used for interaction tests (e.g., by [17, 18]), which consists of including in the model an interaction term to account for interactions between genotype and one environmental covariate at a time:

$$\mathbf{y} = \mathbf{g}\beta_G + \mathbf{E_i}\gamma + \mathbf{g}\mathbf{E_i}\beta_{GxC} + \mathbf{u} + \boldsymbol{\psi}. \tag{46}$$

Here, $\mathbf{E_i}$ denotes a single cellular environment, i.e., one column from matrix $\mathbf{E}$. The parameter $\gamma$ denotes the additive effect of the environment and $\beta_{GxC}$ denotes the effect size of the interaction term. To facilitate a direct comparison, this model employs the same relatedness component as used in CellRegMap ($\mathbf{u}$). Note that using this approach involves performing as many tests as there are cellular contexts ($i = 1, .., n_E$), and thus needs to be followed by multiple testing correction. The results report in Fig. 2 are based on Bonferroni adjustment for the number of environmental variables considered.

#### 4.2 Assessment of statistical calibration and power

We assessed calibration of both CellRegMap and StructLMM under null, assuming no genetic effects (i.e. $\sigma_0^2 = 0$, section 3.5.1) as well as in the case of a persistent genetic effects only (i.e. $\text{var}_{GxC} = 0$, see section 3.5.1). We repeat this simulation study either simulating Gaussian or negative binomial distributed observations (Supplementary Fig. 2.1). In all cases, 200 SNPs were tested and the resulting p-values were compared to values drawn from a uniform distribution to assess calibration (QQ-plots). All analyses were performed both in the case of no relatedness, modelling 2,500 independent cells (one cell per individual) and in the presence of repeated structure (50 cells for each of 50 individuals, resulting in 2,500 cells in total).

For our power analysis, we compared CellRegMap to the SingleEnv-Int model described above, followed by Bonferroni correction across the contexts tested. Power was assessed as the true positive rate across 250 simulated genes. We assessed power between our model and the conventional single-environment interaction test as a function of i) the fraction of genetic variance explained by G×C ($\rho_0$), ii) the number of simulated contexts with G×C and iii) the number of tested cellular environments. We simulated both a fixed number of cells per donor (50 cells per donor, as above), as well as variable number of cells per individual (see section 3.2).

Both for the calibration and power analysis, we either considered the setting of directly simulating Gaussian distributed phenotypes, or alternatively simulating RNA-seq counts by drawing from a negative binomial model (see section 3.5.5); see Fig. 2 and Supp Fig. 2.1 & 2.2. In the case of simulating counts, we used the same preprocessing as employed on real data (quantile normalization of log-transformed counts) to define the input phenotype for CellRegMap and alternative methods.

### 5 Application to endoderm differentiation

We considered single-cell expression data from 125 individuals from [16]. These data cover *in vitro* differentiation of iPS cells from pluripotent stage (day0) to definitive endoderm (day3). Compared to the entire dataset considered in the original publication, we discarded two outlying cell sub-populations (n=475 and n=1,212 respectively).

#### 5.1 Preprocessing of scRNA-seq count data

Count data were processed as in the primary paper [16], where counts were normalised using scran [19] and log-transformed ($log_2(x+1)$). Prior to be inputed into the model as phenotype vectors (i.e., as $\mathbf{y}$), single-cell counts for a given gene were quantile-normalized to better fit the Gaussian distribution assumed by the model. The log-normalized count data (prior to quantile normalization) for the top 500 highly variable genes is also used as input for MOFA (see details below).

#### 5.2 Estimation and annotation of MOFA factors

MOFA [5] factors calculated from the expression profiles of the top 500 highly variable genes identified across all cells (using scran's function "modelGeneVar"), using default parameters. Annotation of the leading 10 MOFA factors was performed using gprofiler [20], considering the absolute loading of individual genes as estimated by MOFA. For each MOFA factor, we considered the top 20 enriched pathways to annotate individual factors (adjusted p-values $< 0.05$; Supplementary Fig. 3.1).

#### 5.3 Testing for G×C effects

We considered *cis*-eQTL from [16]. In particular, we considered all gene-SNP pairs that were significant (FDR $< 10\%$) in one or more of the developmental stages considered in the original paper, i.e., iPSCs, mesendoderm, definitive endoderm. In total, we considered 4,470 eQTL pairs (3,240 eGenes). This approach is typical to GxE interaction testing, where SNPs with at least weak persistent effects (g only) are used as good candidates to display GxE effects.

Next, we mapped context-specific effects using either 1 or 10 MOFA factors. In both cases, all 20 MOFA factors were used to construct the background term, i.e., columns in $\mathbf{E}$ standardized and then used to build $\mathbf{\Sigma} = \mathbf{E}\mathbf{E}^T$. To account for the multiple testing burden, the resulting p-values were corrected at the gene-level using the Bonferroni procedure to control the family-wise error rate, and across genes using the Storey method to control the false-discovery rate (FDR). Significant results were reported at FDR $<$ 5%.

#### 5.4 Estimation of single-cell effect sizes

We estimated both persistent genetic effects ($\beta_G$) and cell-level effect sizes due to G×C ($\boldsymbol{\beta_{GxC}}$; as described in Section 2) for the top significant context-specific eQTL identified (FDR $< 5\%$), when considering either only one MOFA factor (MOFA 1, capturing differentiation) or the first 10 MOFA factors as cell contexts.

### 6 Application to neuronal differentiation

Next, we considered single-cell transcriptomic data from over 200 individuals from [21]. We focused on a single cell type: midbrain dopaminergic neurons (DA), across three conditions defined in the original publication: day 30, day 52 untreated, and day 52 rotenone-treated. In total, this consists of 135,435 cells across 210 donors.

#### 6.1 Preprocessing of scRNA-seq count data

Count data were processed as in the primary paper [21], where counts were normalised using scanpy [22] and log-transformed ($log_2(x+1)$). Prior to be inputed into the model as phenotype vectors (i.e., as $\mathbf{y}$), single-cell counts for a given gene were quantile-normalized to better fit the Gaussian distribution assumed by the model.

#### 6.2 Pseudo-cell calculation

We obtained pseudo-cells by clustering similar cells in UMAP space using an approach similar to approaches described in [23, 24]. Briefly, for each cell type we calculated the first 50 PCs across all conditions and donors, using all expressed genes (after QC, n=32,738). Next, we applied batch-correction for the pool id using Harmony [25], as implemented in scanpy ('scanpy.external.harmony_integrate'). The Harmony-corrected PCs were then used to build a k-NN (k=10) graph based on Euclidean distances, separately for each condition and donor. Subsequently, cells were clustered using scanpy's implementation of the Leiden algorithm ([26], as implemented in 'scanpy.tl.leiden') at a resolution of 3.4.

The pseudo-cell calculation described above resulted, in these data, in a total of 8,479 pseudocells (10-40 pseudocells per donor, 10-100 cells per pseudocell).

#### 6.3 Clustering of single-cell effect sizes

Next, we set out to cluster context-specific eQTL based on their allelic effects due to G×C. First, we considered 212 eQTL which displayed significant G×C effects using our method (FDR $<$ 5%) when considering the leading 10 MOFA factors as cellular contexts. Next, we considered the estimated effect size profiles due to G×C (as in Section 2). These profiles were normalised to contain only positive values between 0 and 1. Finally, clustering on the normalised profiles was performed using SpatialDE [27], by using the same first 10 MOFA factors as spatial coordinates, and default parameters. This identified 12 clusters, containing between 2 and 43 genes (main text Fig. 4b).

##### 6.4 Cluster enrichment

For each cluster, we considered Pearson's correlation between the cluster's summary profiles (as outputted by SpatialDE) and single-cell gene expression across all genes (n=32,738). Next, we selected all genes with positive correlation larger than 0.4 and used gprofiler [20] to identify enriched pathways (considering Gene Ontology (GO) biological processes, molecular function, cellular components, pathways from KEGG Reactome and WikiPathways; miRNA targets from miRTarBase and regulatory motif matches from TRANSFAC; tissue specificity from Human Protein Atlas; protein complexes from CORUM and human disease phenotypes from Human Phenotype Ontology). The gprofiler function ("g:GOSt" as implemented in R) performs over-representation analysis on input gene list (ordered by correlation level) using a hypergeometric test, corrected for multiple testing. The latter is performed using a tailor-made method (g:SCS algorithm) which analytically approximates a threshold t corresponding to the 5% upper quantile of randomly generated queries of the provided size. All actual p-values resulting from the query are transformed to corrected p-values by multiplying these to the ratio of the approximate threshold t and the initial experiment-wide threshold a=0.05.). Finally, we selected the top 3 significant (adjusted p-values $< 0.05$) pathways per cluster (Supplementary Fig. 4.2).

### 7 Derivations

#### 7.1 Detailed derivations from section 2

A linear mixed model is a type of Gaussian Process, that can be written as:

$$\mathbf{f} \sim \mathcal{GP}\{\mathbf{m}; k(\mathbf{X}, \mathbf{X})_{\boldsymbol{\theta}}\}, \tag{47}$$

or, equivalently, as:

$$\mathbf{f} = \mathbf{m} + \mathbf{u}, \tag{48}$$

where

$$\mathbf{m} = \mathbf{X}\boldsymbol{\beta}_{\theta} \text{ and } \mathbf{u} \sim \mathcal{N}(\mathbf{0}, k(\mathbf{X}, \mathbf{X})_{\theta}). \tag{49}$$

When we model our data ($\mathbf{X}$ and $\mathbf{y}$), we consider $\mathbf{y}$ as being a sample from the random variable $\mathbf{f}$, and $\mathbf{X}$ as being fixed. Let us collectively call the parameters $\boldsymbol{\theta}$. For all parameters, we can find the set of best linear unbiased predictor (BLUP) estimators ($\hat{\boldsymbol{\theta}}_{BLUP}$ or simply $\hat{\boldsymbol{\theta}}$) which i) minimise the variance and ii) are unbiased [14]. We identify $\hat{\boldsymbol{\theta}}$ by maximising the likelihood:

$$\hat{\boldsymbol{\theta}} = \text{argmax}\{p(\mathbf{y})_{\boldsymbol{\theta}}\} \tag{50}$$

Let us reason about out-of-sample $\mathbf{f}_*$, assuming that we now have the BLUP-estimated parameters ($\hat{\boldsymbol{\theta}}$):

$$\begin{bmatrix} \mathbf{f} \\ \mathbf{f}_* \end{bmatrix} = \mathcal{N}\left( \begin{bmatrix} \mathbf{m}_{\hat{\boldsymbol{\theta}}} \\ \mathbf{m}_{*\hat{\boldsymbol{\theta}}} \end{bmatrix}; \begin{bmatrix} k(\mathbf{X}, \mathbf{X})_{\hat{\boldsymbol{\theta}}} & k(\mathbf{X}, \mathbf{X}_*)_{\hat{\boldsymbol{\theta}}} \\ k(\mathbf{X}_*, \mathbf{X})_{\hat{\boldsymbol{\theta}}} & k(\mathbf{X}_*, \mathbf{X}_*)_{\hat{\boldsymbol{\theta}}} \end{bmatrix} \right). \tag{51}$$

We want to know $\mathbf{y}_*$, it seems reasonable (argue better, maybe mentioning that this is also a BLUP estimation) to define (noise-free prediction according to [15])

$$\mathbf{y}_* := \mathrm{E}[\mathbf{f}_* | \mathbf{y}]_{\hat{\boldsymbol{\theta}}}. \tag{52}$$

We can show [15] that (note that we omit $\hat{\boldsymbol{\theta}}$ for reading purposes):

$$\mathbf{f}_*|\mathbf{f} \sim \mathcal{N}\left(\mathbf{m}_* + k(\mathbf{X}_*, \mathbf{X})k(\mathbf{X}, \mathbf{X})^{-1}(\mathbf{f} - \mathbf{m});\ k(\mathbf{X}_*, \mathbf{X}_*) - k(\mathbf{X}_*, \mathbf{X})k(\mathbf{X}, \mathbf{X})^{-1}k(\mathbf{X}, \mathbf{X}_*)\right). \tag{53}$$

Therefore (needs some steps to prove),

$$\mathbf{y}_* = \mathbf{m}_* + k(\mathbf{X}_*, \mathbf{X})k(\mathbf{X}, \mathbf{X})^{-1}(\mathbf{y} - \mathbf{m}). \tag{54}$$

In our case,

$$\mathbf{m} = \mathbf{W}\boldsymbol{\alpha} + \mathbf{g}\beta_G + \mathbf{E}\boldsymbol{\gamma}, \tag{55}$$

$$k(\mathbf{X}, \mathbf{X}) = \mathbf{K} = \sigma^2_{GxE}(\mathbf{g} \odot \mathbf{E})(\mathbf{g} \odot \mathbf{E})^T + \sigma^2_{re}\mathbf{R} \odot \mathbf{E}\mathbf{E}^T + \sigma^2_n\mathbf{I}. \tag{56}$$

Note that

$$k(\mathbf{X}_*, \mathbf{X}) = \mathbf{K}_{(*,.)} = \sigma^2_{GxE}(\mathbf{g}_* \odot \mathbf{E}_*)(\mathbf{g} \odot \mathbf{E})^T + \sigma^2_{re}\mathbf{R}_{(*,.)} \odot \mathbf{E}_*\mathbf{E}^T + \sigma^2_n\delta(\mathbf{X}_*, \mathbf{X}), \tag{57}$$

where $\delta$ is the Dirac distribution (0 when $i \neq j$ and 1 when $i = j$).

Therefore,

$$\mathbf{y}_* = \mathbf{W}_*\boldsymbol{\alpha} + \mathbf{g}_*\beta_G + \mathbf{E}_*\boldsymbol{\gamma} + k(\mathbf{X}_*, \mathbf{X})k(\mathbf{X}, \mathbf{X})^{-1}(\mathbf{y} - \mathbf{W}\boldsymbol{\alpha} + \mathbf{g}\beta_G + \mathbf{E}\boldsymbol{\gamma}) \tag{58}$$

Now, for each individual, we consider its environmental profile $\mathbf{e}_{*,i}$ (the $i^{th}$ corresponding row from $\mathbf{E}_*$), and estimate $y_{*,i}(\text{ref})$ when setting $g_{*,i} = 0$ and then $y_{*,i}(\text{alt})$ when setting $g_{*,i} = 1$ and obtain the corresponding allelic effect $\beta^*_{GxE,i}$ as the difference, normalised by a constant (based on p=MAF):

$$\beta^*_i = \frac{1}{\sqrt{2p(1-p)}}\ (y_{*,i}(\text{alt}) - y_{*,i}(\text{ref})) \tag{59}$$

For all samples, we consider the vectorial form:

$$\boldsymbol{\beta}^* = \frac{1}{\sqrt{2p(1-p)}}\ (\mathbf{y}_*(\text{alt}) - \mathbf{y}_*(\text{ref})), \tag{60}$$

where:

$$\mathbf{y}_*(\text{alt}) = \mathbf{y}_*(\mathbf{g}_* = \mathbf{1}) = \mathbf{m}_*(\mathbf{g}_* = \mathbf{1}) + \mathbf{K}_{(*,.)}(\mathbf{g}_* = \mathbf{1})\ \mathbf{K}^{-1}(\mathbf{y} - \mathbf{m}) \tag{61}$$

and

$$\mathbf{y}_*(\text{ref}) = \mathbf{y}_*(\mathbf{g}_* = \mathbf{0}) = \mathbf{m}_*(\mathbf{g}_* = \mathbf{0}) + \mathbf{K}_{(*,.)}(\mathbf{g}_* = \mathbf{0})\ \mathbf{K}^{-1}(\mathbf{y} - \mathbf{m}). \tag{62}$$

Let us set:

$$\mathbf{v} = \mathbf{K}^{-1}(\mathbf{y} - \mathbf{m}). \tag{63}$$

Next, we note that:

$$\mathbf{m}_*(\mathbf{g}_* = \mathbf{1}) - \mathbf{m}_*(\mathbf{g}_* = \mathbf{0}) = \mathbf{W}_*\boldsymbol{\alpha} + \mathbf{1}\beta_G + \mathbf{E}_*\boldsymbol{\gamma} - (\mathbf{W}_*\boldsymbol{\alpha} + \mathbf{E}_*\boldsymbol{\gamma}) = \mathbf{1}\beta_G. \tag{64}$$

Moreover,

$$\begin{aligned} \mathbf{K}_{(*,.)}(\mathbf{g}_* = \mathbf{1}) - \mathbf{K}_{(*,.)}(\mathbf{g}_* = \mathbf{0}) = \\ \sigma^2_{GxE}\mathbf{E}_*(\mathbf{g} \odot \mathbf{E})^T + \sigma^2_{re}\mathbf{R}_{(*,.)} \odot \mathbf{E}_*\mathbf{E}^T + \sigma^2_n\mathbf{I} + \\ -(\sigma^2_{re}\mathbf{R}_{(*,.)} \odot \mathbf{E}_*\mathbf{E}^T + \sigma^2_n\mathbf{I}) = \\ \sigma^2_{GxE}\mathbf{E}_*(\mathbf{g} \odot \mathbf{E})^T \end{aligned} \tag{65}$$

Thus,

$$\mathbf{y}_*(\text{alt}) - \mathbf{y}_*(\text{ref}) = \mathbf{1}\beta_G + \sigma^2_{GxE}\mathbf{E}_*(\mathbf{g} \odot \mathbf{E})^T\mathbf{v} \tag{66}$$

And finally, by substituting eq. (66) in eq. (60):

$$\boldsymbol{\beta}^* = \frac{1}{\sqrt{2p(1-p)}}\ \{\underbrace{\mathbf{1}\beta_G}_{\boldsymbol{\beta_{G^*}}} + \underbrace{\sigma^2_{GxE}\mathbf{E}_*(\mathbf{g} \odot \mathbf{E})^T\mathbf{K}^{-1}(\mathbf{y} - \mathbf{W}\boldsymbol{\alpha} - \mathbf{g}\beta_G - \mathbf{E}\boldsymbol{\gamma})}_{\boldsymbol{\beta_{GxE^*}}}\}, \tag{67}$$

where $\mathbf{K}$ is as defined in eq. (56).

#### 7.2 Detailed derivations from section 3

Consider eq. (33) for a single individual:

$$y = y_0 + \sum_i^S g_i \beta_{G,i} + \sum_i^S g_i \boldsymbol{\epsilon}^T \boldsymbol{\alpha}_{GxE,i} + \boldsymbol{\epsilon}^T \boldsymbol{\gamma} + \tag{68}$$

and let us focus on the GxE term:

$$y_{GxE} = \sum_i^S g_i \boldsymbol{\epsilon}^T \boldsymbol{\alpha}_{GxE,i} \tag{69}$$

We have

$$\mathrm{E}[y_{GxE}] = \mathrm{E}[\sum_i^S g_i \boldsymbol{\epsilon}^T \boldsymbol{\alpha}_{GxE,i}] = \sum_i^S \mathrm{E}[g_i \boldsymbol{\epsilon}^T \boldsymbol{\alpha}_{GxE,i}] = \sum_i^S \overbrace{\mathrm{E}[g_i]}^{=0} [\boldsymbol{\epsilon}^T \boldsymbol{\alpha}_{GxE,i}] = 0, \tag{70}$$

where we use that $\mathbf{g}_i$ is standardised ($\mathrm{E}[g_i] = 0$) and we assume that $g_i$ and $\boldsymbol{\epsilon}^T \boldsymbol{\alpha}_{GxE,i}$ are uncorrelated.

We also have (for every SNP $i$, environment $j$):

$$\mathrm{E}[y_{GxE}^2] = \mathrm{E}[(\sum_i^S g_i \boldsymbol{\epsilon}^T \boldsymbol{\alpha}_{GxE,i})^2] = \sum_j \mathrm{E}[\epsilon_j^2]\ \mathrm{E}[\alpha_{i,j}^2] = \sigma_i^2, \tag{71}$$

after a couple of assumptions. We define $\sigma_i^2 = v_i$ if $\mathbf{g}_i$ is causal and $\sigma_i^2 = 0$ otherwise. We assume all causal SNPs to have equal effect as defined by $v_i = \sigma^2 / n_{GxE}$, where $n_{GxE}$ is the number of SNPs having GxE effects. We also assume that $\mathrm{E}[\epsilon_j] = 0$ and $\mathrm{E}[\epsilon_j^2] = \frac{1}{n_E}$ for every environment $j$.

#### 8 Software availability

CellRegMap is available under an open source license at `https://github.com/limix/CellRegMap`. Code to reproduce the specific analyses presented can be accessed under `https://github.com/annacuomo/CellRegMap_analyses`.

[20] Uku Raudvere, Liis Kolberg, Ivan Kuzmin, Tambet Arak, Priit Adler, Hedi Peterson, and Jaak Vilo. g: Profiler: a web server for functional enrichment analysis and conversions of gene lists (2019 update). *Nucleic acids research*, 47(W1):W191–W198, 2019.
